# High-Throughput Phenotyping of Seed Quality Traits Using Imaging and Deep Learning in Dry Pea

**DOI:** 10.1101/2024.03.05.583564

**Authors:** Mario Andres Morales, Hannah Worral, Lisa Piche, Atanda Sikiru Adeniyi, Francoise Dariva, Catalina Ramos, Khang Hoang, Changhui Yan, Paulo Flores, Nonoy Bandillo

## Abstract

Seed traits, such as seed color and seed size, directly impact seed quality, affecting the marketability and value of dry peas [1]. Assessing seed quality is integral to a plant breeding programs to ensure optimal seed standards. This research introduced a phenotyping tool to assess seed quality traits specifically tailored for pulse crops, which integrates image processing with cutting-edge deep learning models. The proposed method is designed for automation, seamlessly processing a sequence of images while minimizing human intervention. The pipeline standardized red-green-blue (RGB) images captured from a color light box and used deep learning models to segment and detect seed features. Our method extracted up to 86 distinct seed characteristics, ranging from basic size metrics to intricate texture details and color nuances. Compared to traditional methods, our pipeline demonstrated a 95 percent similarity in seed quality assessment and increased time efficiency (from 2 weeks to 30 minutes for processing time). Specifically, we observed an improvement in the accuracy of seed trait identification by simply using an RGB value instead of a categorical, non-standard description, which allowed for an increase in the range of detectable seed quality characteristics. By integrating conventional image processing techniques with foundational deep learning models, this approach emerges as a pivotal instrument in pulse breeding programs, guaranteeing the maintenance of superior seed quality standards.

## 1 Introduction

Dry pea (Pisum sativum L., 2n = 14) belongs to the Leguminosae (or Fabaceae) family, is the second most important grain legume in the world after common bean (Phaseolus vulgaris L.) and is considered one of the most genetically diverse crop species. [2–4]. It is currently grown in most temperate regions of the world for human consumption, livestock feed, and the pet food market [5]. It is capable of fixing atmospheric nitrogen through association with specialized soil bacteria called rhizobia that act as a natural fertilizer, thus greatly reducing the dependency on synthetic fertilizers [6–9].

Dry pea is also a valuable source of dietary proteins, mineral nutrients, complex starch and fibers [10], and its nutritional profile complements that of cereals [11]. To date, dry pea remains a nongenetically modified organism (non-GMO), which is appealing to consumers, and has contributed to this crop emerging as a leader in the plant-based protein product market. Beyond Meat has released a burger that contains approximately 20 grams of pea protein per burger [12], and Ripple has created a niche in the market for non-dairy milk with its Ripptein-made pea milk [13]. As the demand for dry pea products increases, it is important to further improve the yield and quality of the seed to meet these needs [14].

The quality of pulse crop seed is an important determining factor for the marketability and value of the crop in both domestic and global plant markets [15]. The quality of dry pea seed is mainly judged by its visual appearance, which includes the size, shape, and color of the seed. According to the Dry Pea and Lentil Council of the US, these criteria remain primary requirements to obtain a premium price in the current market for dry pea. Furthermore, seed quality traits tend to be associated with seedling vigor [16], and a significant positive correlation has been observed between seed size and grain yield [17]. Thus, the pulse crop breeding program routinely evaluates seed quality traits such as seed size, color, and shape to ensure the marketability of any cultivar released through the program.

The conventional method of evaluating seed quality in dry pea requires visual inspection of color and sieve analysis to estimate seed size and 100 seeds weight. This method can be subjective, laborious, and time-consuming, while generating a limited amount of data to be used in making an effective selection. Therefore, the conventional method of assessing quality can take months when screening over a thousand new breeding lines, creating a selection bottleneck in the breeding pipeline. The development of a rapid and accurate evaluation tool to collect accurate, interpretable, and biologically meaningful seed quality traits on a large scale is key to supporting rapid breeding decisions to meet consumer preferences.

Computer vision techniques have emerged as a promising alternative to manual methods in addressing this phenotyping bottleneck. Numerous academic and industrial approaches have used image analysis and machine learning to characterize seed traits. However, many current methodologies require extensive calibration for precise seed segmentation or positioning [18–35], and some advanced industrial solutions like the VideometerLab [36], and the advanced platform for executing such analyses, can be cost prohibitive for public breeding programs.

To address these phenotyping challenges, our goal was to develop an accessible high-throughput seed phenotyping pipeline suitable for pulse crop breeding applications. This open source method combines image processing with deep learning models to enable automated large-scale seed phenotyping. By standardizing image capture conditions and using deep learning for robust seed detection and trait quantification, our pipeline can significantly increase the accuracy and efficiency of phenotyping compared to manual techniques. This document provides a detailed overview of the end-to-end pipeline that we created to alleviate the phenotyping bottleneck [37] and accelerate genetic gain for critical consumer preferences, such as seed quality traits in pulse crops.

## 2 Materials and Methods

### 2.1 Plant Materials

In total, 287 USDA germplasm accessions (including 191 from the Pea Single Plant Plus Collection PSP [38]) were used for data capture and pipeline design. This germplasm represents the different regions and taxonomies of the species and reflects the morphological diversity of the seed. Accessions were collected from 64 geographical origins and are now kept and distributed by the Western Regional Plant Introduction Station, USDA, Agricultural Research Service (ARS), Pullman, WA.

### 2.2 Field Experiments

All NDSU lines were planted in an augmented row-column design with five repeated checks [39, 40] during the 2021 and 2022 growing seasons at the North Dakota Agricultural Experiment Station in Prosper (47’00’N, 97’11’W) and the North Central Research Extension Center in Minot (27’29’N, 109’56’W), North Dakota, United States. Seeds were treated with fungicide and insecticide before planting. At planting, 30 seeds were planted on 152 × 60 cm plot size with 30 cm spacing between the plots. The plots were harvested at physiological maturity (90-120 days after planting) and dried to a moisture content of 15 percent.

### 2.3 Color Light Box Capturing Building

While commercial color light boxes [41, 42] are available, most lack an integrated and stable structure to position a phone camera directly above the imaging area. Furthermore, existing commercial solutions can be prohibitively expensive for multi-research station use. Consequently, we developed an affordable Do It Yourself (DIY) light box design optimized for top-down seed imaging using widely accessible materials [43]. The Color Light Box was constructed using a standard 18×18×18inch cardboard box sourced from common shipping stores [44], which provides an enclosed imaging area painted matte black to minimize light reflections. A Pringles potato chip can cap [45, 46] was modified into a Petri dish-shaped insert 4 inch in diameter to hold seed samples. This insert was fixed adjacent to a United States quarter coin glued to the box floor, providing a size reference in the images. The phone camera was mounted on top of the box. Detailed building instructions are provided in the Supplementary Materials (Appendix 1). The basic construction materials used are inexpensive and available internationally, and the quarter coin size reference allows adaptation with local coinage of similar dimensions where needed. Alternative boxes have been developed using a similar approach but require more tools and effort [47].

### 2.4 Imaging pre-processing and General Pipeline Overview Description

After setting up a color lightbox as described in Appendix 1, we captured high-resolution images of seeds. These images were taken under controlled lighting conditions to ensure consistency. We used a dark, non-reflective background to enhance the contrast between the seeds and their surroundings, making it easier to analyze the images (see Figure 1A). To prepare the images for analysis, we used an algorithm from the Pliman R package. This tool helped us in several ways: it located a reference object (the coin) in the image, rotated the image to a standard orientation, and adjusted for any distortions caused by the camera lens or variations in color. This was important to make sure that all images were consistent and comparable (see Figure 1B). Initially, we experimented with traditional image processing methods using OpenCV [48] and Pliman [49], but we found these methods to be lacking. They required many manual adjustments to correctly identify and segment (or separate) objects in the images, which was time consuming and prone to errors. In spring 2023, our approach took a significant turn with the release of Meta AI’s Segment Anything Model (SAM) [50], a highly advanced tool trained on over a billion examples. SAM’s Blue Box, depicted in Figure 2, was a game changer for us. Its strength lies in its ability to automatically and accurately segment various objects in an image, including the seeds of interest. This meant that SAM could identify and isolate each seed from the rest of the image with high precision. The final step in our process was to train a model called YOLOv8 [51] to automatically recognize and label seeds in new images. YOLOv8 is an object detection model designed to identify specific objects within an image and understand their characteristics. By training YOLOv8 with the images we had prepared and processed through SAM, we could automate the task of annotating new images with high accuracy (see Figure 1D).

**Figure 1:**
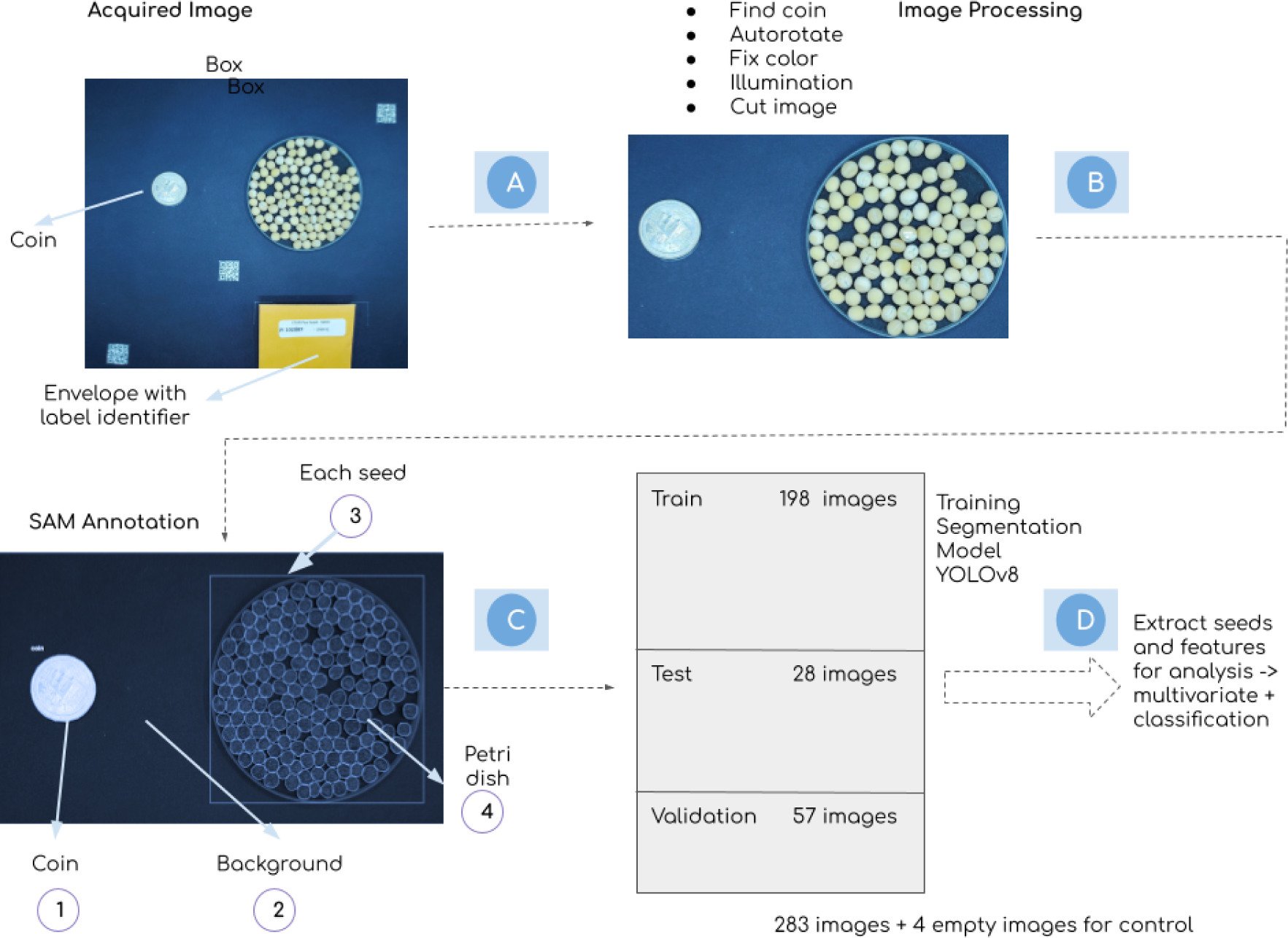
Image Processing Pipeline for Image Processing, Annotation and Segmentation for Dry Pea images: (a) Image Acquisition; (b) Image Processing; (c) SAM Annotation and (d) YOLOv8 Segmentation Model

**Figure 2:**
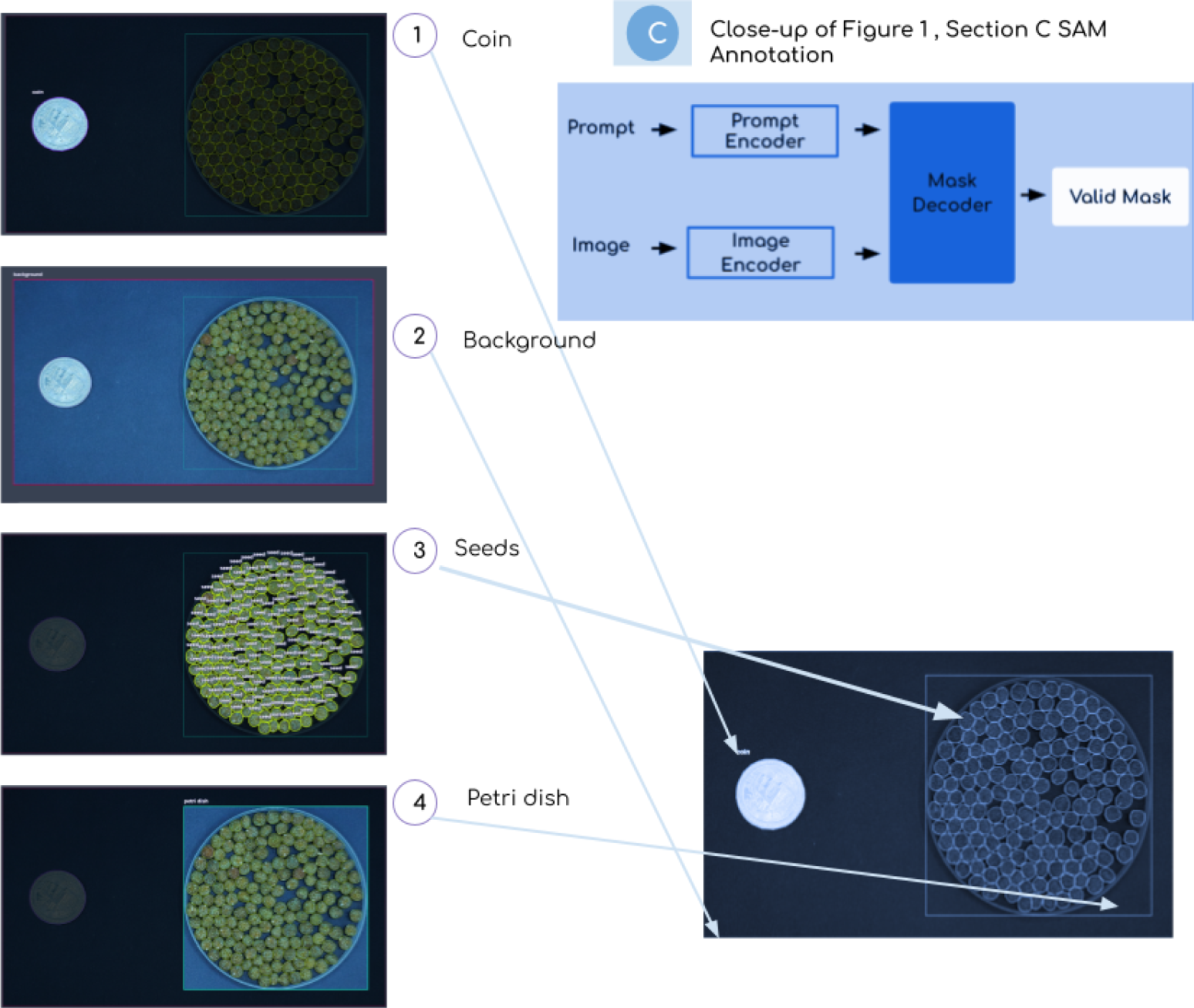
”Featured is a detailed close-up of the Segment Anything Model (SAM) Annotation Model. Central to this display is the “Blue Box”, which contains an in-depth description of the model’s architecture. This architecture is crucial in understanding how SAM operates to distinguish and annotate various objects in an image. Within this Blue Box, the SAM model is delineated into four distinct classes for annotation, each designed to recognize and categorize different objects within an image. These classes are: 1) Coin: This class is trained to identify coins in the image. The coin serves as a reference object, aiding in the standardization of the image scale and providing a basis for comparative analysis. 2) Background: The background class focuses on distinguishing the non-essential parts of the image, effectively separating the background from the main objects of interest. This helps isolate the seeds, the Petri dish, and the coin from their surroundings, ensuring a more precise segmentation. 3) Seeds: This class is dedicated to identifying and annotating seeds. It accurately pinpoints each seed, regardless of its position or overlap with other seeds, allowing a detailed analysis of individual seed characteristics. 4) Petri dish: The Petri dish class is tasked with recognizing and delineating the Petri dish in the image. This is crucial for context, as it helps to establish the spatial relationship between the seeds and their container. Each class plays a vital role in the comprehensive analysis and annotation process, contributing to the ability of the SAM model to accurately segment and identify various objects within a single image.

### 2.5 Seeds Segmentation using Segment Anything Model SAM

A well-annotated data set is crucial before training an object detection model such as YOLOv8 [51]. For seeds, this process involves precise segmentation to distinguish individual seeds and accurate annotation to provide the model with the necessary information about the location of each seed. This was achieved using the Segment Anything Model (SAM) of Meta AI [50] for efficient and accurate seed segmentation. SAM is a foundation model for segmentation, trained on a vast dataset (more than a billion masked annotations), making it capable of segmenting a wide range of objects, including seeds and coins. Its architecture comprises three primary modules: an image encoder, a prompt encoder, and a mask decoder (see Figure2 Blue Box).

SAM processes the images and returns segmentation masks, effectively separating the seeds from the background. For each segmented seed, a bounding box is drawn around it defined by its top left and bottom right coordinates. This rectangular box encapsulates the seed, providing the spatial information necessary for YOLOv8 (You Only Look Once) training. The output is formatted as JSON, which was chosen for its compatibility and ease of integration with our training process. In particular, the decision to exclude data augmentation from our methodology was informed by the controlled conditions within our lightbox setup and the consistent positioning of the seeds, ensuring uniformity and precision in our data collection.

### 2.6 Seeds Detection using Yolov8

After labeling our seed images, we trained them using a deep learning model based on the YOLO version 8 architecture. YOLO models are known for their speed and accuracy in detecting objects, and they work by analyzing an entire image in one go, unlike traditional methods that scan parts of an image separately. For our study, we used Python’s Tensorflow 2 [52] library to run YOLOv8 on our specially annotated data set. This involved tasks such as segmenting and detecting specific objects in new images. YOLO architecture [53] takes the entire image and divides it into a grid of squares of equal sizes (SxS grid).

Each square in this grid is like a mini version of the whole picture. By breaking the image into smaller areas, YOLO can focus on specific parts of the image one at a time, making it easier and faster to find and recognize objects. This is like looking closely at each small section of a map to understand what is there, rather than trying to take in the whole map at once. Each cell of the grid predicts multiple bounding boxes and their associated class probabilities. For seed detection, the classes would be different types of seeds or seed-related anomalies. The input layer typically takes an image of size 416×416×3 (height, width, channels).

The YOLO architecture uses several convolutional layers to extract features from the input image [53] (see Figure 3). In our case, these layers capture spatial hierarchies and patterns, essential for distinguishing between different types of seed, the coin, the background, and the Petri dish. Each grid cell in the detection layer predicts two key metrics: first, the bounding box coordinates that determine the object location and size within the image and second, the confidence score that reflects the model level of certainty regarding the presence of an object in the box and accuracy of prediction. Additionally, the YOLO architecture incorporates class probabilities that determine the likelihood that the detected object belongs to a particular class. To reduce the number of boxes that overlap, YOLO uses a technique known as ‘nonmax suppression’. This technique retains the bounding box with the highest confidence score while suppressing other boxes with significant overlap. The YOLO architecture further employs a multipart loss function that takes into account: Classification loss (pertaining to the detection of an object in the box), localization loss (relating to the accuracy of the bounding box coordinates), and confidence loss (relating to boxes with and without objects). The backbone of YOLOv8 is built on the CSPDarknet53 architecture, which includes 53 convolutional layers. These layers are designed to extract features from the images, allowing the model to identify subtle differences within the seeds. The Head section of YOLOv8 consists of multiple convolutional layers followed by fully connected layers. The primary function of a convolutional layer is to predict the bounding boxes and associated probabilities. The inclusion of a self-attention mechanism allows the model to assign varying levels of importance to different features in the seed images (see Figure 3 for a visual representation of the described operation and architecture).

**Figure 3:**
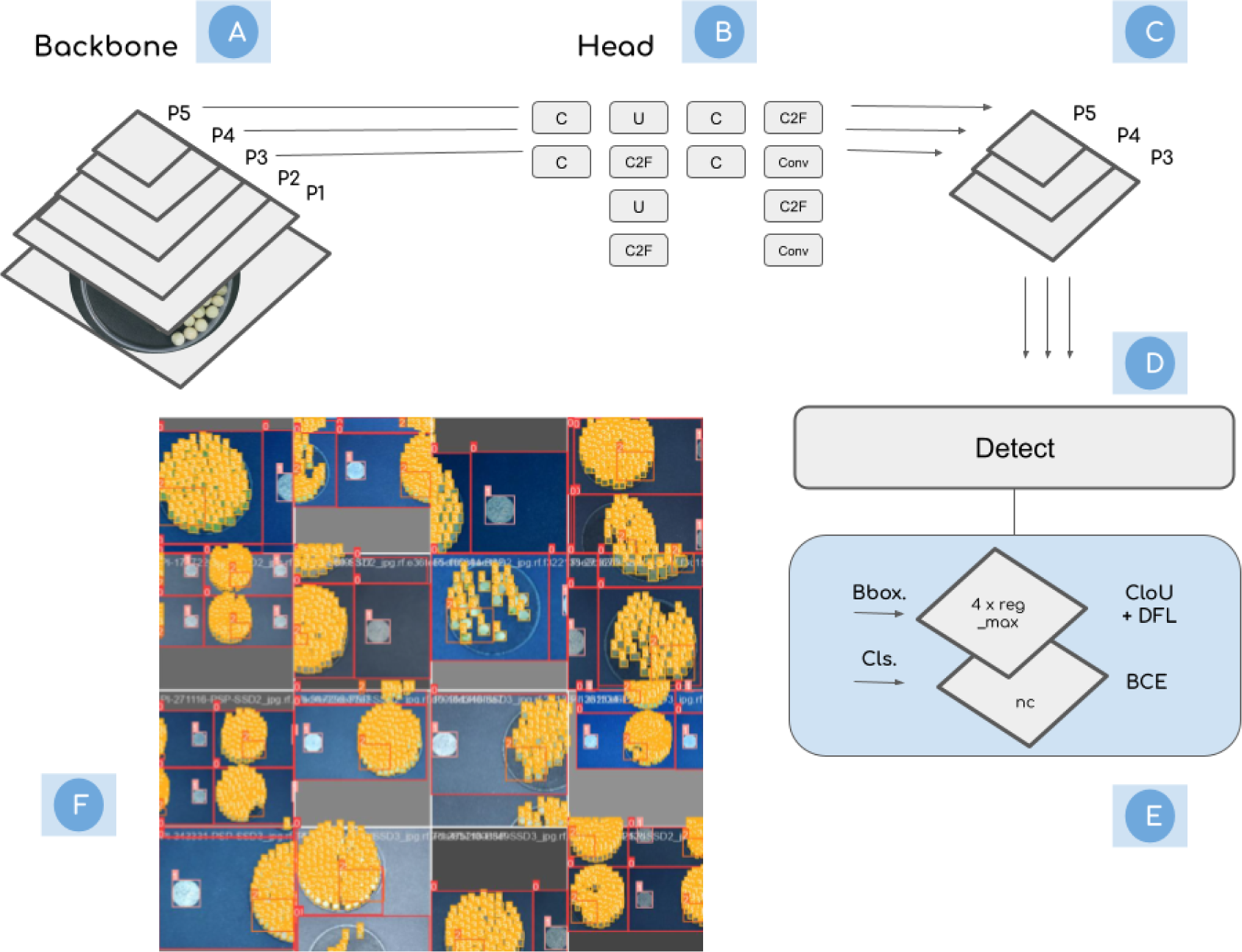
The YOLOv8 architecture, tailored for seed anomaly detection, operates on an input image of dimensions 416×416×3. Its backbone (a), CSPDarknet53, consists of 53 convolutional layers designed to extract intricate features from images, enabling the model to discern subtle variations in seeds. The head section (b) integrates multiple convolutional layers followed by fully connected layers, primarily predicting bounding boxes and their associated probabilities. Each cell in the detection layer (c,d) provides bounding box coordinates and a confidence score. To minimize overlapping boxes, YOLO employs ‘nonmax suppression’ (e). Additionally, the architecture computes class probabilities and incorporates a multi-part loss function. A unique self-attention mechanism within the model assigns varied importance levels to different characteristics of the seed image, enhancing its detection capabilities. (f) show some of the convolutional operations on the image

We conducted two sets of experiments with our model, one with 50 epochs and the other with 300, to see how well the model learned over time. Our results showed that running 50 epochs was sufficient for our needs. In these experiments, YOLOv8 was trained to recognize four different objects: the coin, the background, the seeds and the Petri dish. Our dataset for this included 198 training images, 28 for testing, and 57 for validation. It is worth noting that four of these images were controls that did not contain seeds. To address concerns about whether our data set was large enough to train and evaluate the YOLOv8 model, we considered the following: •YOLOv8 Efficiency: This particular model is effective at learning from smaller datasets thanks to its unique approach of analyzing images in one pass. This means that it can generalize from limited data more effectively.

- Data quality and germplasm diversity: Although our data set was not large, it was diverse and contained a wide range of seed types (accessions). This variety, spread over 198 training images, 28 tests, and 57 validation images, allowed for thorough learning and application to new data.
- Controlled Experiment Design: Our experiments, particularly the one with 50 epochs, showed that the model could learn effectively from the dataset we provided.
- Cross-Validation Methodology: We used cross-validation in our study, which is a technique to maximize the use of a dataset for training. It involves rotating different subsets of data for training and evaluation, improving the accuracy and reliability of the model.
- Inclusion of Control Images: By including images without seeds, we improved the model’s ability to accurately identify the presence of seeds.

Although a larger dataset could improve the performance of the model, our current dataset was sufficient for the objectives of our study. Future research could build on this foundation, using more data to potentially further improve the model performance.

### 2.7 Infrastructure and Hyperparameters Tuning

In our experimental setup, we used two distinct computational resources. Nvidia V100 GPU, as detailed by Yang et al. [54] was sourced from Google Colaboratory [55] and was instrumental for the 50-epoch run. In contrast, for the more extensive 300-epoch run, we leveraged the Nvidia A100 (2nd Gen) GPU [56] available at the Local Cluster CCAST [57]. During the training phase of the YoloV8 model, we opted for the Adam optimizer, renowned for its superior stochastic optimization capabilities. A critical parameter, the learning rate, which dictates the extent of weight adjustments during the training process, was set at 0.001. Our training regimen processed samples in groups, with each batch containing 64 samples. Several specific parameters were integral to the model’s architecture and functioning. The model was designed to accept input images with dimensions of 608 x 608 pixels. For object detection, it employed three unique shapes of bounding boxes, called anchors. A detection is deemed valid when its associated confidence score exceeds the 0.25 mark. To manage overlapping bounding boxes, we implemented a Non-Max Suppression (NMS) mechanism with a threshold set at 0.45. Additionally, the degree of overlap between bounding boxes was ascertained using an intersection-over-union (IOU) metric, with a threshold value of 0.5. Delving deeper into the model architecture, each convolutional layer was equipped with 64 filters. In total, the model encompassed 53 such convolutional layers. We also incorporated the LeakyReLU activation function to introduce nonlinearity into the model.

### 2.8 Feature Engineering

In our study, we identified 86 unique traits to describe germplasm. These traits can be subsequently employed for inferential genomics in subsequent laboratory analyses.

The features span several domains:

**Color** (15 features): These metrics include the mean (mean), spread (standard deviation), and asymmetry (skewness) of the color intensities for each BGR (Blue-Green-Red) channel of the image. In simpler terms, the mean represents the average color intensity, the standard deviation gauges the range of color intensities, and the skewness measures the tilt of the color distribution.

**Shape** (10 features): This category includes geometric properties [58, 59] of the seed such as its area, perimeter, convexity, solidity, minor and major axis lengths, orientation, eccentricity, circularity, and aspect ratio. The area measures the seed size, while the perimeter defines its boundary length. Convexity assesses the alignment of the seed’s boundary with its convex outline, and solidity compares the area of the seed to that of its convex outline.

**Texture** (15 features):

Texture Base Metrics (2 features): These metrics include skewness and kurtosis.

Haralick Texture Descriptors[60, 61] (13 features): Derived from a Gray Level Cooccurrence Matrix (GLCM), these descriptors capture the texture characteristics, patterns, and variations in each seed image’s appearance. The GLCM is essentially a matrix that represents the frequency of occurrence of pixel pairs with specific gray levels at a particular spatial relationship. From this matrix, various statistical measures are extracted, such as the Angular Second Moment (indicating image uniformity), contrast (measuring local variations), correlation (evaluating linear dependency of pixel pairs), and several others. Each of these measures provides information about different aspects of the image’s texture.

**Elliptical Fourier Descriptors**[58, 59] (EFD) (20 features): These descriptors offer a precise way to describe the contours of seed images. They are based on the contour coordinates of the seed image and allow the decomposition of a contour into a series of ellipses. This decomposition aids in analyzing and classifying the shapes of seeds according to their morphological characteristics.

**LBP histogram**[62] (26 features): The Local Binary Pattern Histogram Texture Descriptors (LBP) is a set of 26 texture descriptors that characterize the spatial structure of local image textures. The LBP method involves comparing each pixel in the image with its neighbors and encoding the result as a binary number. The histogram of these numbers then serves as a texture descriptor, useful for texture classification. In essence, the LBP histogram provides a way to analyze the texture properties of the image by accumulating the occurrences of each LBP code in the image.

### 2.9 Statistical Analysis

To evaluate the precision and robustness of a deep learning framework to extract seed traits from pulse crops, specifically dry peas, the primary objective was to observe the relationship between manual ground-truth observations from 67 accessions (measured), focusing on the lengths of the major and minor axes and measurements computed using image processing. Linear regression and correlation analysis explored the potential relationships between the observed seed sizes and those derived by the model and the other extracted descriptors. Additionally, the average colors extracted from the images were visually verified to ensure accuracy. The potential descriptors identified through these analyses will be used in future genomic research.

## 3 Results

### 3.1 SAM and Yolov8 Model’s Performance

The Segment Anything Model (SAM) demonstrated a notable proficiency in segmentation and labeling tasks within our dataset, achieving a threshold that surpasses the rate of 96.11 percent. This performance is indicative of SAM’s robust capabilities in accurately segmenting and identifying distinct elements within the dataset. Further evaluation of the YOLOv8 model was performed using our seed data set from Pisum sativum L. The model achieved a precision metric of 0.977, reflecting its high precision in correctly identifying and classifying seeds. This level of precision is significant, particularly considering the potential applications of YOLOv8 in automated analysis for agricultural quality control, specifically seeds. The findings of this study suggest that YOLOv8 holds promise as a reliable model for seed analysis and classification tasks. The successful application of this model in our research could serve as a basis for future investigations into the use of advanced object detection models in agricultural studies. In particular, the ability of YOLOv8 to extract and utilize diverse features could be instrumental in advancing research in seed diversity and related fields. (see Figures 4, 5). In reference to Figure 6, it’s important to clarify the process and outcomes related to the detection of the Petri dish in our study. Initially, our model attempted to detect the entire Petri dish during the extraction phase, but it encountered challenges in achieving high precision. This led us to explore an alternative method aimed at enhancing the detection accuracy. However, even with this modified approach, the precision in detecting the entire Petri dish did not meet our desired threshold. Consequently, after careful consideration, we decided to exclude the Petri dish class from our final model analysis. This decision was made to maintain the overall accuracy and reliability of our results, acknowledging that the detection of the Petri dish did not achieve the high precision we aimed for in our study.

**Figure 4:**
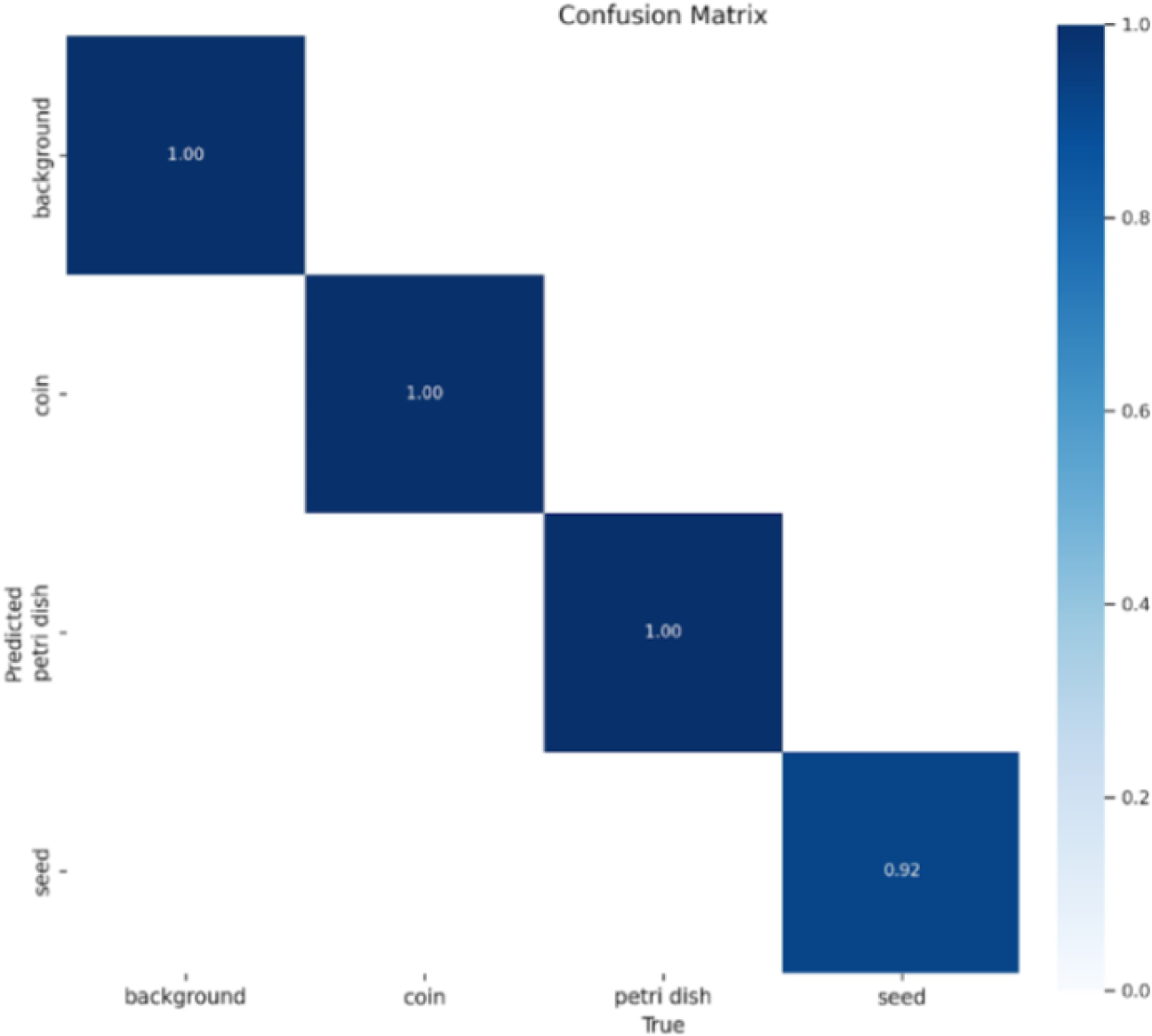
Performance of Yolov8 on Pisum sativum L. accessions dataset: Confusion Matrix with classification classes, background, coin, Petri dish and seeds

**Figure 5:**
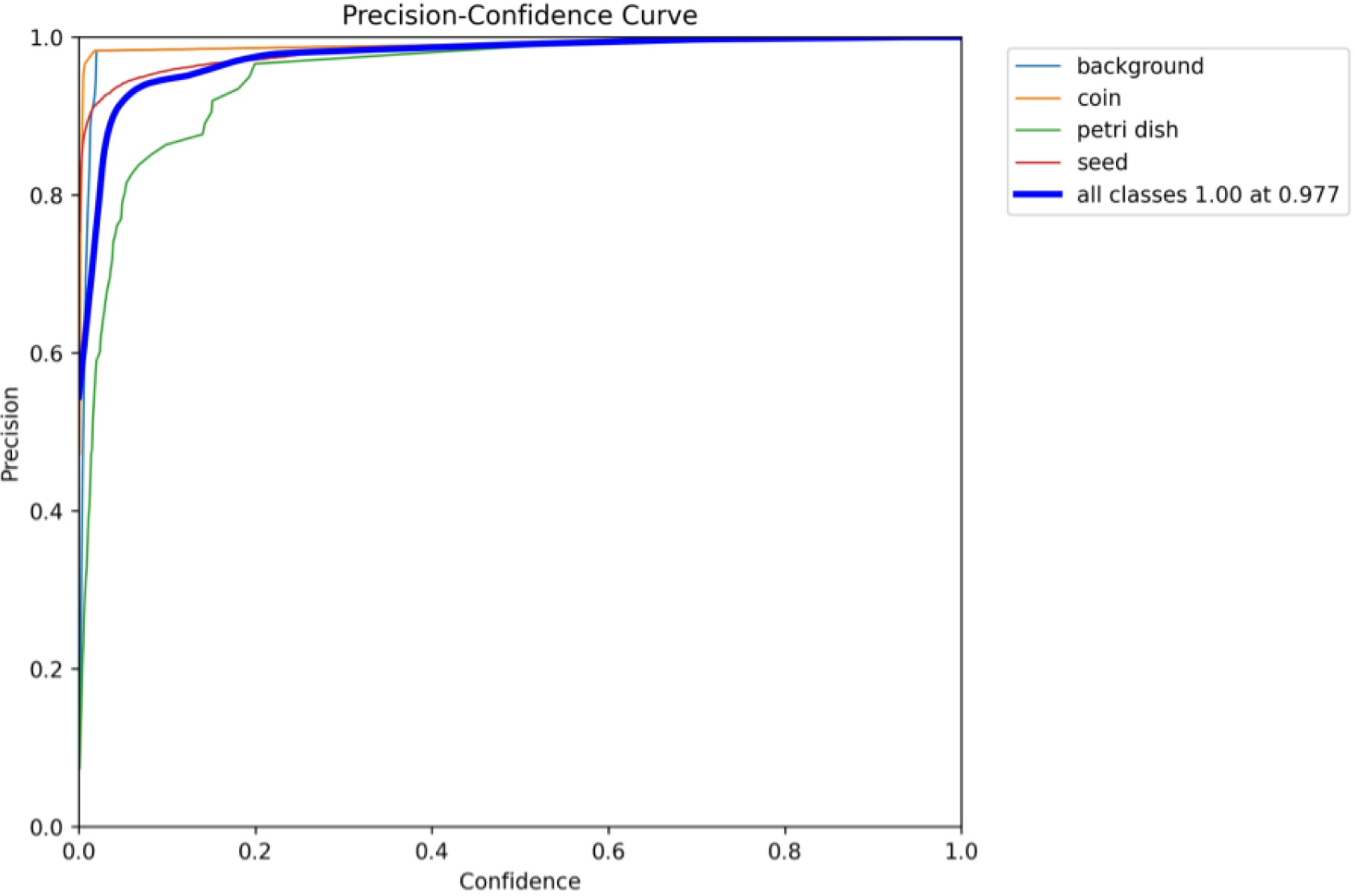
AUC performance curve with precision per class and overall 0.977.

**Figure 6:**
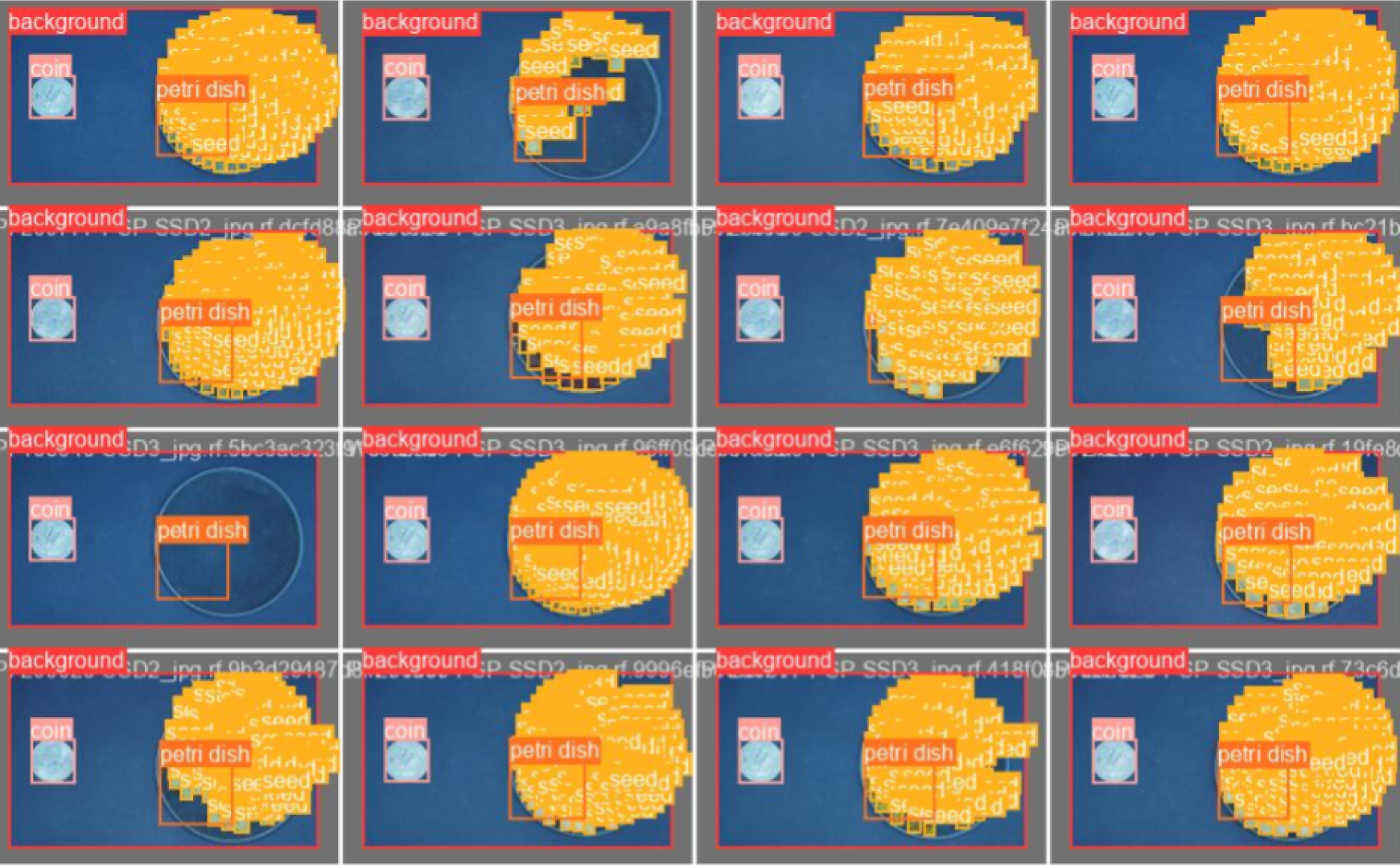
Validation Results with Petri dish detected but not covering the whole region, these are different accessions

An important observation is that the detection of the Petri dish class is not achieved with high precision on the whole dish, so we wanted to use it for Region of Interest (ROI) detection; however, we can remove the class for simplicity. See Figure 6.

### 3.2 Model Ground Truth Validation

In our study, we conducted size comparisons between manual observations and measurements derived from a deep learning model on 67 dry pea accessions. For each accession, two seeds were meticulously measured for their major and minor axis lengths using a caliper. These seeds were then subjected to analysis via a deep learning model to determine the lengths of their major and minor axes.

A notable correlation coefficient of 0.95 emerged between the manual measurements and those obtained from the model. This robust correlation underscores the effectiveness of the deep learning approach in accurately extracting seed traits from image data.

We used linear regression to further investigate the relationship between the manually observed seed sizes and those determined by image processing. The analysis yielded an *R*^2^ value of 0.85, indicating that the measurements obtained from image processing can account for 85 percent of the variability in the observed seed sizes. The closeness of this value to 1 reinforces the deep learning model’s proficiency in mirroring manual measurements of seed traits.

The combined evidence from the high correlation coefficient and the value of *R*^2^ accentuates the potential of the deep learning framework as an efficient tool for the extraction of high-throughput seed traits, especially in pulse crops. Although our focus in this study was on the major and minor axes, the model demonstrated the ability to extract other seed descriptors. Future studies will delve into these descriptors in a deeper sense, as preliminary correlation analyzes have identified several promising seed trait indicators. Detailed summaries of the regression models can be found in Figure 7 and Appendix 2.

**Figure 7:**
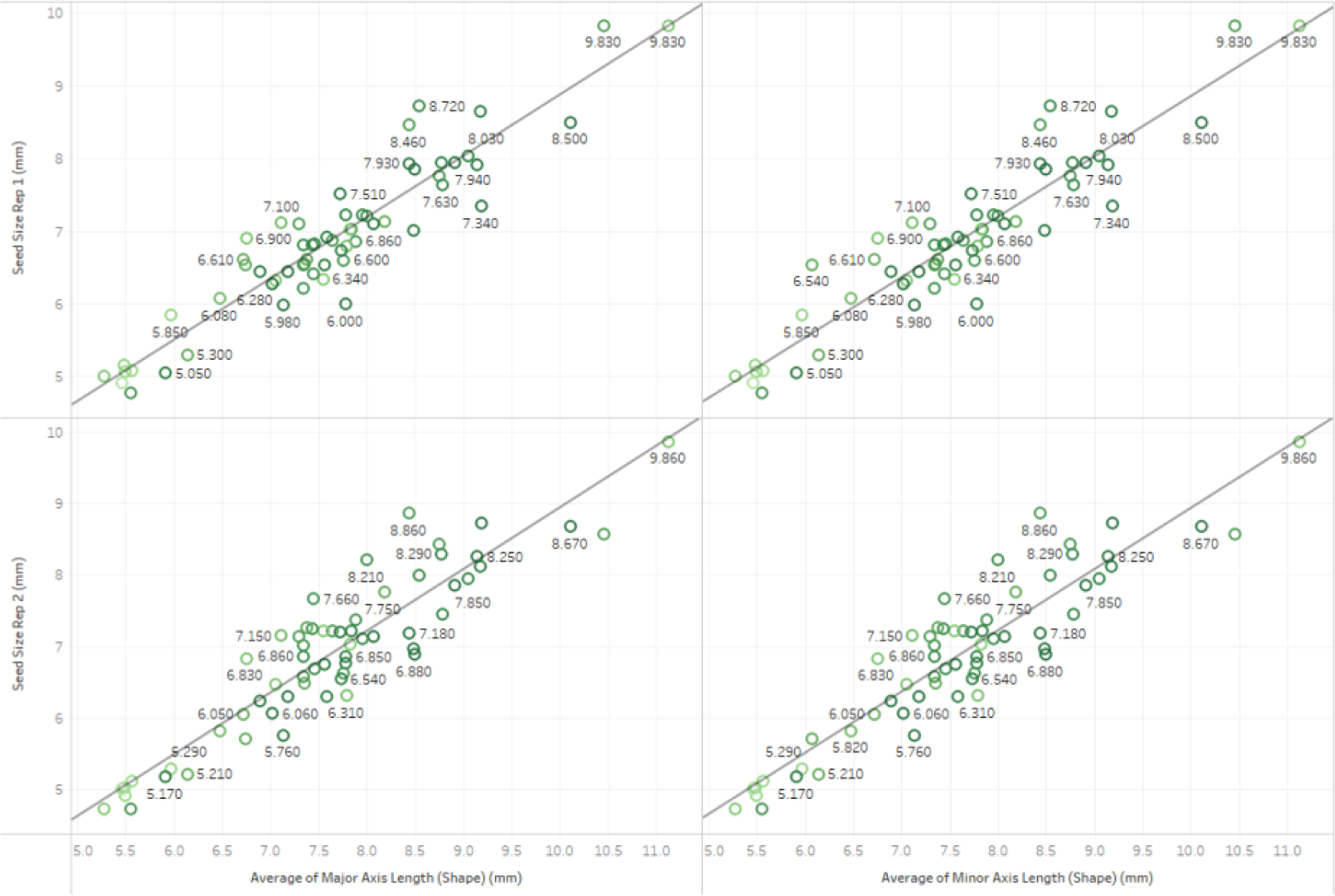
Linear Regression Comparison for Seed Size Validation Results, see appendix 2 for statistical summaries

### 3.2.1 Color comparison

In our analysis, we focused on comparing the colors between the original images and those extracted from individual seeds, as depicted in Figure 8. Although our standard methodology for color comparison is primarily categorical, limiting our ability to validate the colors statistically, we employed a qualitative approach to ensure color accuracy. To validate the color consistency, we conducted a detailed visual examination in which experts compared the color hues, saturation, and brightness between the original images and the extracted colors. This figure is crucial as it visually demonstrates the effectiveness of our color extraction technique, highlighting the fidelity of the colors extracted from the seeds compared to their original counterparts. The primary objective of including this figure and analysis in our model was to ensure that the color representation of the seeds is accurate and consistent, which is vital for the precise identification and classification in our study. By demonstrating the color equivalence, we aim to reinforce the reliability of our model in handling color-sensitive tasks, which is essential in applications where color accuracy is paramount, such as in quality control or phenotypic analysis in agricultural studies.

**Figure 8:**
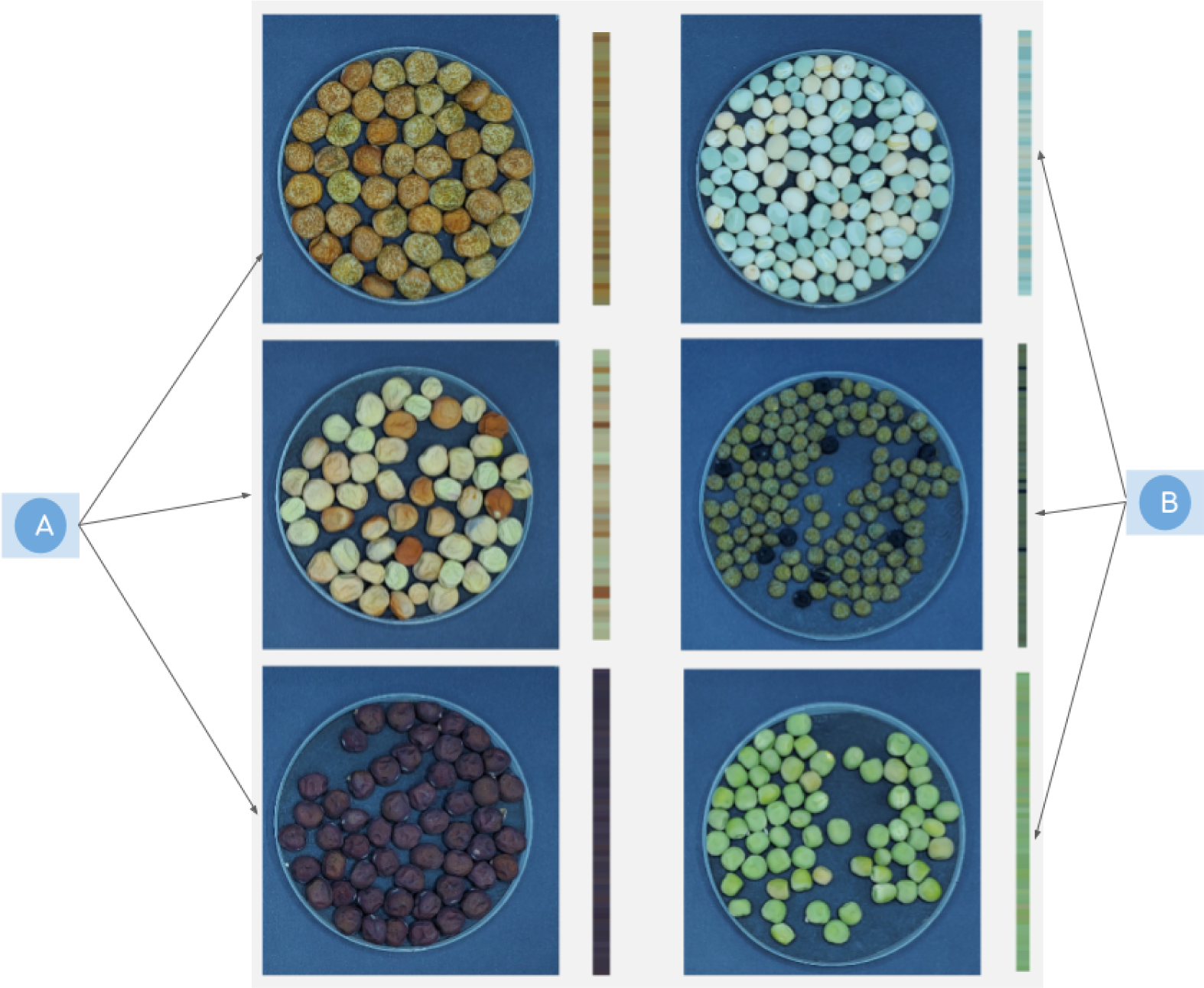
Color Validation Results, (a) Original Images and (b) Histograms of Red Blue Green Colors for each seed captured from the images

## 4 Discussion

The development and implementation of a pipeline for automated analysis of seed traits, especially through deep learning models, presents a multitude of advantages, including efficiency, scalability, data integration, accuracy, and a controlled environment over more conventional methods. Automated processes, by design, tend to reduce manual intervention, which can lead to faster results, especially when analyzing large datasets. Unlike manual measurements, which may carry individual biases, an automated pipeline ensures data processing and measurement extraction consistency. As data sets grow or the need for increased resolution arises, automated pipelines can be easily scaled to accommodate larger data volumes without proportionally increasing time or labor resources. Automated systems often allow seamless integration with other digital platforms or datasets, paving the way for more holistic analyses. The value of *R*^2^ achieved in our regression model not only indicates the robustness of the model, but also emphasizes the precision of our deep learning approach. This high coefficient of determination suggests that the extracted image characteristics provide a reliable representation of the true traits of the seeds. Further strengthening this observation is the statistical significance of the model, as evidenced by a p-value of less than 0.05. This ensures confidence in deploying the model for practical applications and extrapolating its results. Operating in a controlled environment minimizes external variability, ensuring that the data obtained are predominantly influenced by the subject (in this case, seed traits) rather than extraneous factors. This improves the reliability of the results and ensures that the observed variations are due to intrinsic traits of the seeds and not environmental factors.

Our setup currently does not include a barcode system, which is a limitation. Barcodes are essential to provide unique identifiers for individual samples or batches, facilitating easy tracking, data retrieval, and integration. Without it, there is a potential risk of mislabeling or data mismatch, especially when handling large data sets. In addition, although the model performs well under controlled conditions, its efficacy in diverse real-world settings (for example, in the field with a cell-phone application) needs validation. Model refinement might be needed when introduced to varied environments and multiple backgrounds, and additional deep learning models are needed to account for these problems. In conclusion, while our deep learning framework offers promising results in seed trait extraction, continual refinement and addressing the existing limitations will further enhance its utility and application breadth.

## Acknowledgments

Author Contributions

MM and NB designed the investigation. JS and LP manually processed the images prior to sending them to the automated pipeline. LP, KS, and EH captured the images in the DIY lightbox. MM designed and built the box and developed the methods. MM and CR developed and executed the code locally and in the cluster. HW, LP, AR, GR, KS, EH, ASA, RAS, JK, JPJ, SRA, HN, KS, EH and MM processed the plant material before image capture. MM, CR, SA, and NB wrote the initial manuscript with the help of CR in the Appendix for the box construction document and Bibtex preparation. All authors contributed to the corrections and the final version. KH supported the research on the use and deployment of code on the CCAST GPU nodes under his guidance. CY provided algorithmic guidance.

## Data Availability

Scripts to conduct analyzes and directions to build a DIY color lightbox are available on the following github repository.

https://github.com/Bandillo-Lab/hopeseed

## Supplementary Materials

Supplementary Materials will be available in our Github repository and include:

Supplementary File Appendix 1: Box construction

Supplementary file Appendix 2: Regression analysis output of Avg seed size ground truth and Avg minor-axis lengths extracted from the images by accession.

Supplementary file Appendix 3: Correlation Matrix Ground Truth Values and Major and Avg Minor Axis Lengths extracted from the images by accession.

