## Supplementary Materials for "High-Throughput Phenotyping of Seed Quality Traits Using Imaging and Deep Learning in Dry Pea"

**Supplementary File**

**Appendix 1 Box Construction**

Materials List:

1. Box

- Box Provider UPS 8X8X8 cm and UPS cm 10X10X10
- Box thickness 42 mm
- UPS is in over 220 countries and territories from the US and Puerto Rico

<https://www.ups.com/in/en/support/international-tools-resources/country-specific-guidelines-regulations.page>

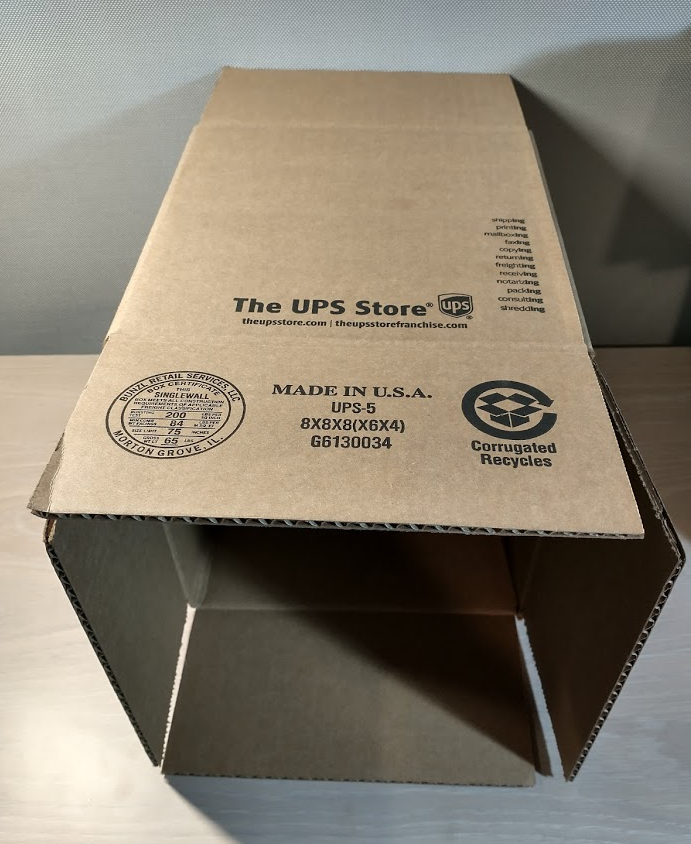

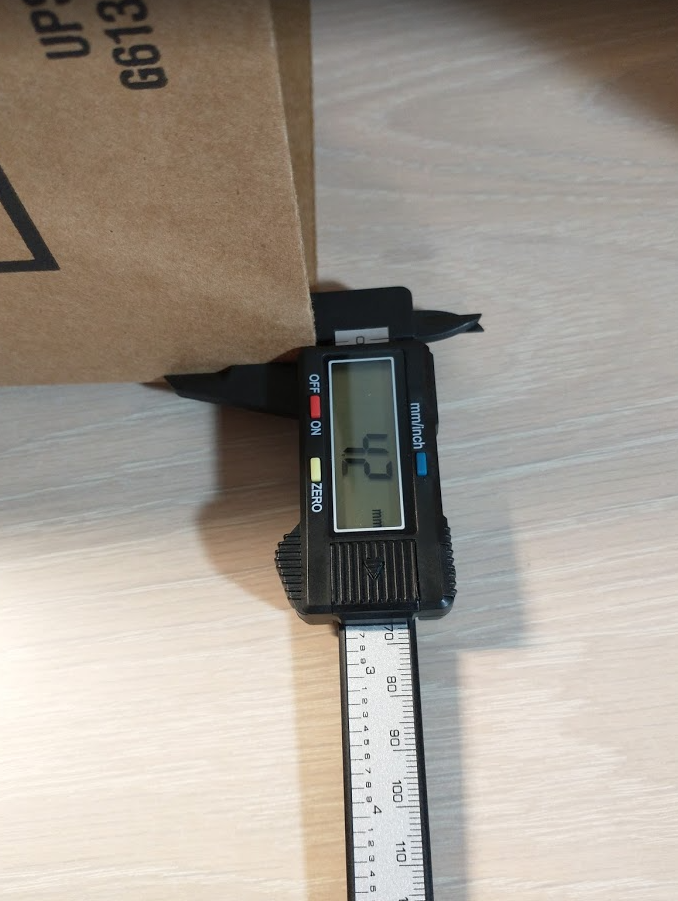

2. Electronic digital caliper

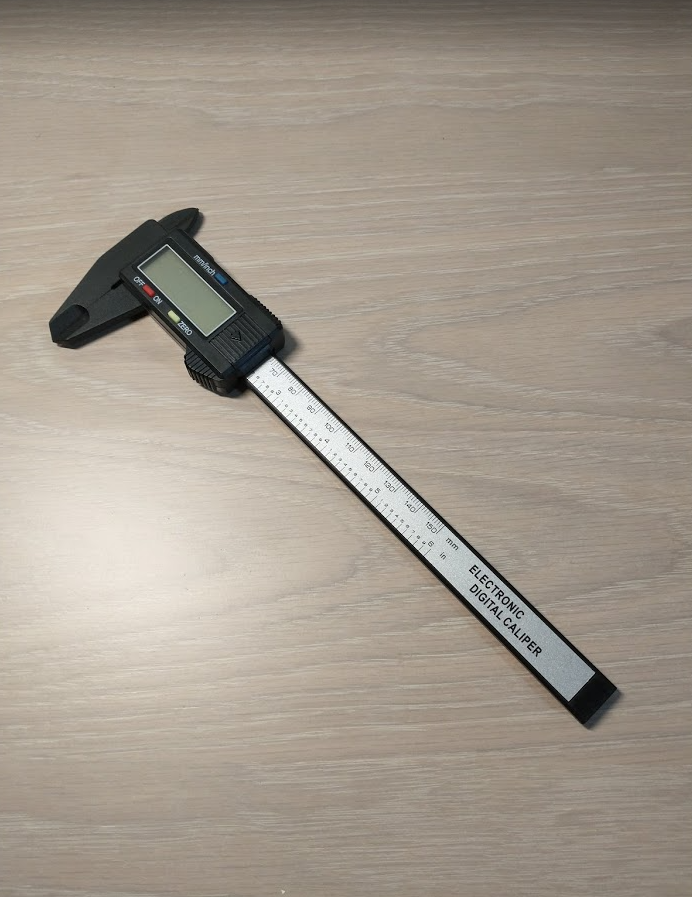

3. Velcro Brand - Sticky Back Hook And Loop Fasteners, Cut-To-Length, 7/8", 12 Count

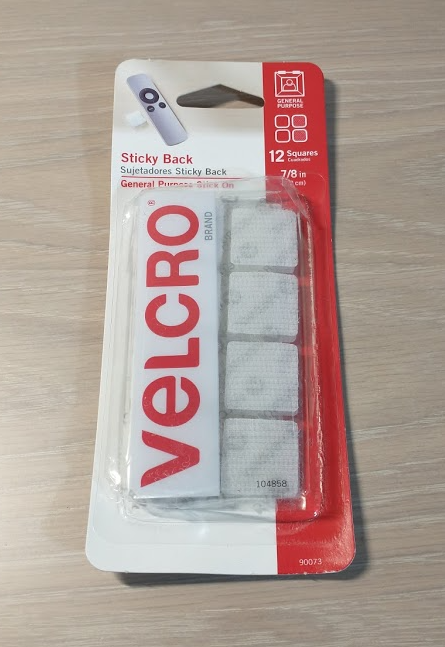

4. LED Lights

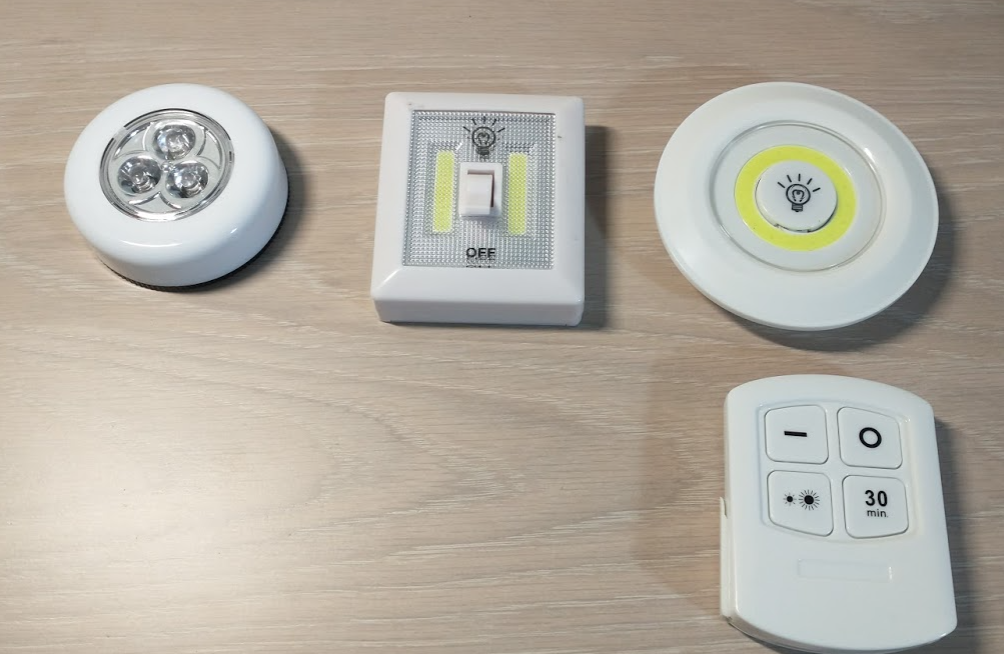

5. Aluminum foil paper

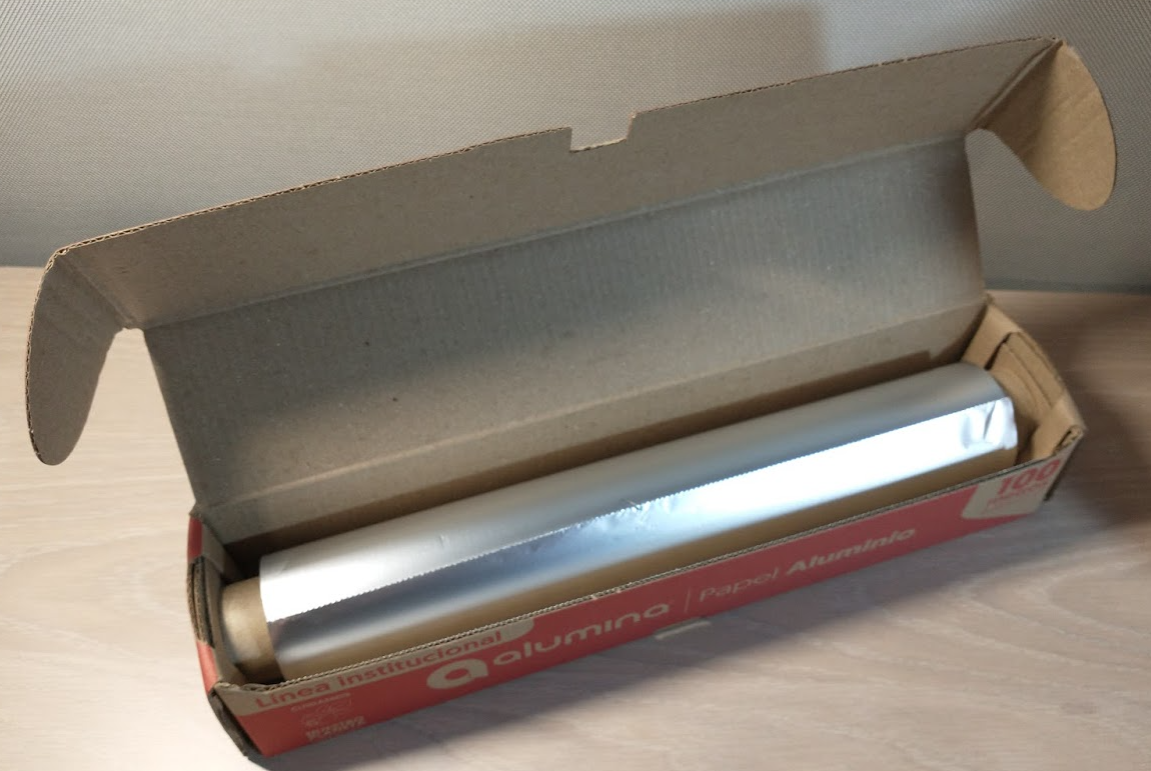

6. Plastic lid of a Pringles 1.42 Oz package

- Pringles are available in over 140 countries

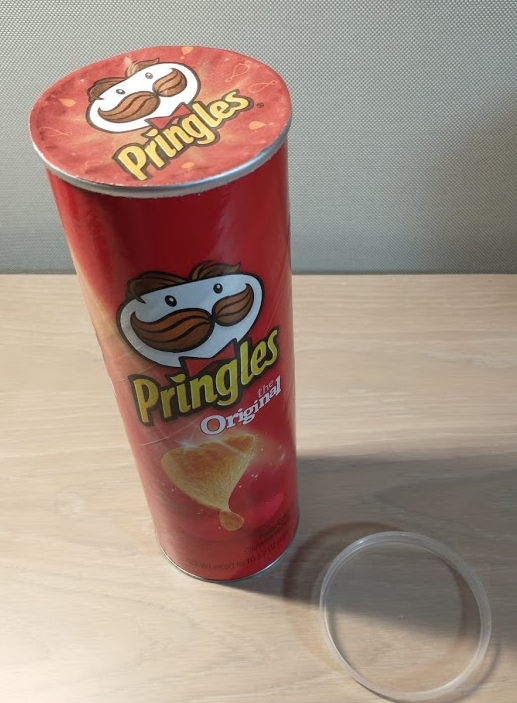

- Lid outside measurements: 77.8 mm

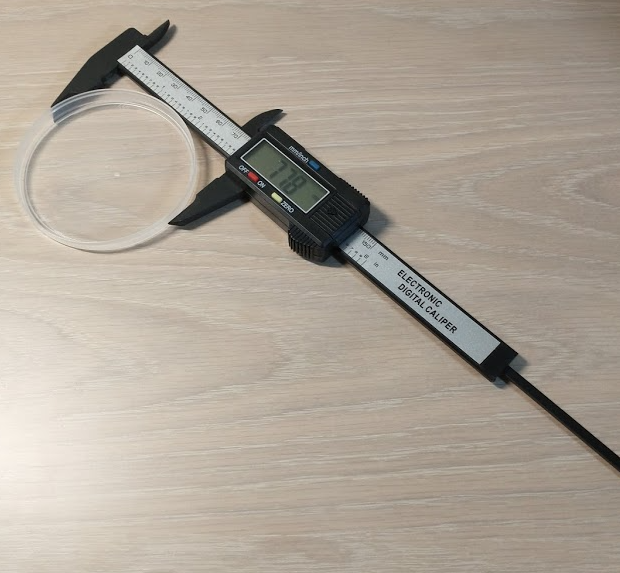

- Lid inside measurements: 76.2 mm

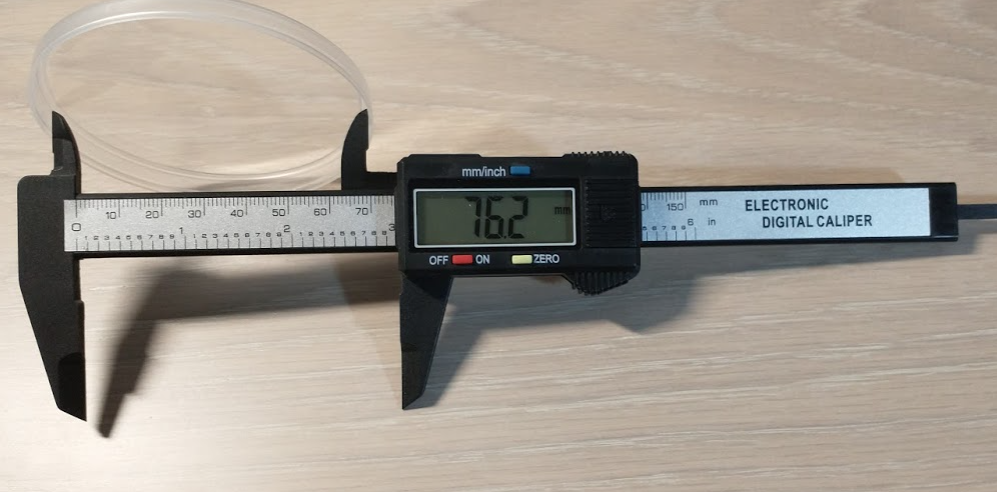

7. Pen compass

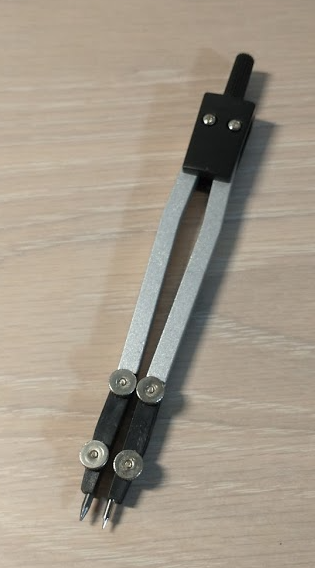

8. Tesa glue pen

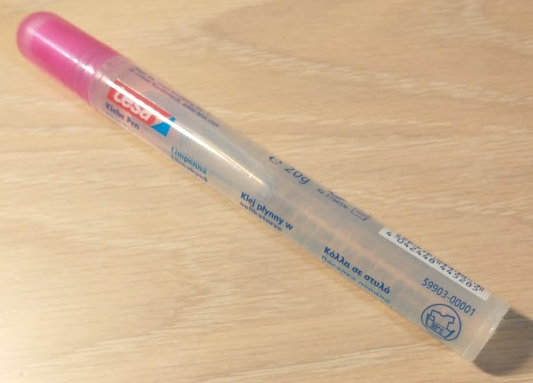

9. Black cardboard

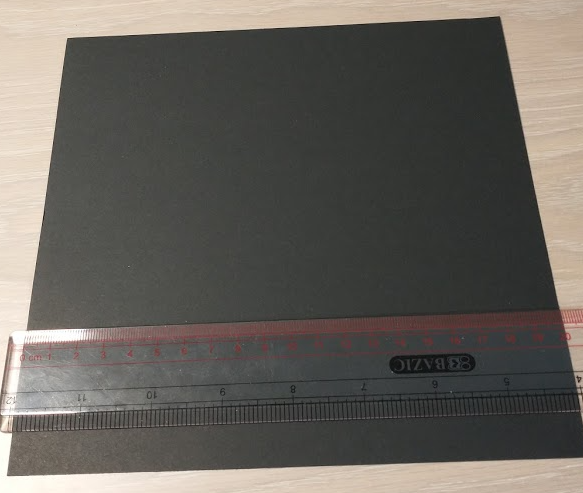

10. 1 USD quarter dollar or equivalent

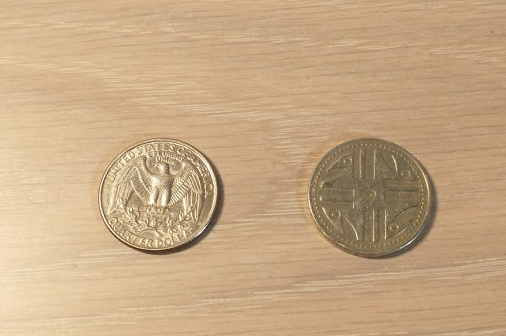

11. Plastic ruler

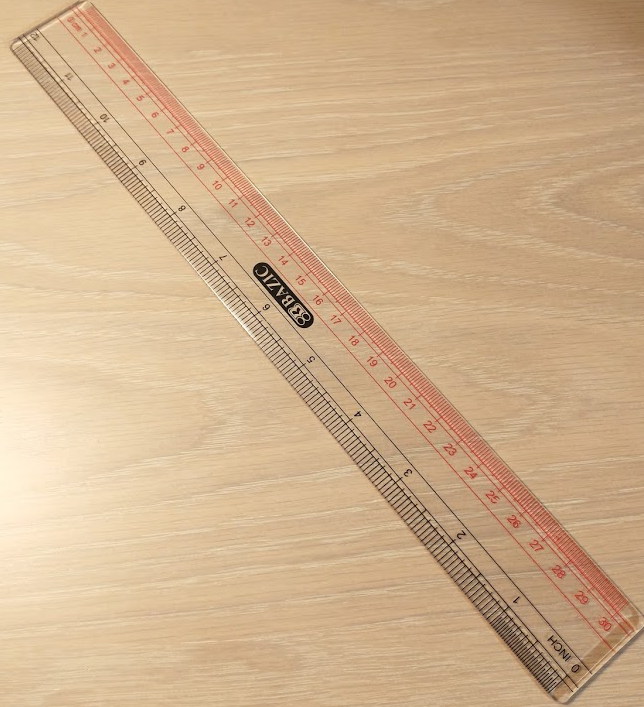

12. Scalpel

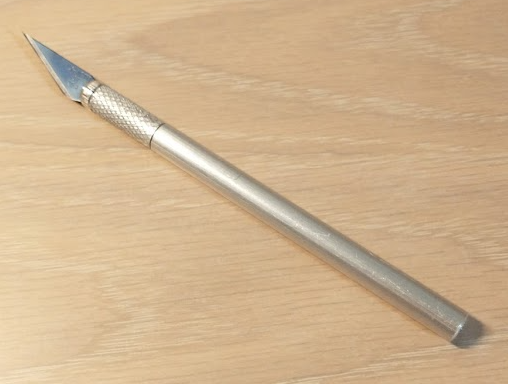

13. Scissors

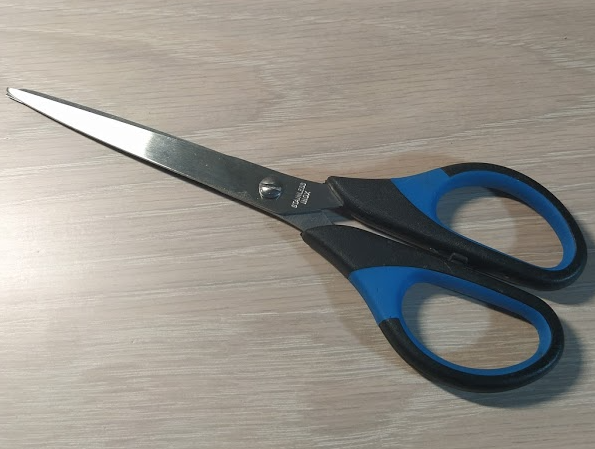

14. Black pen

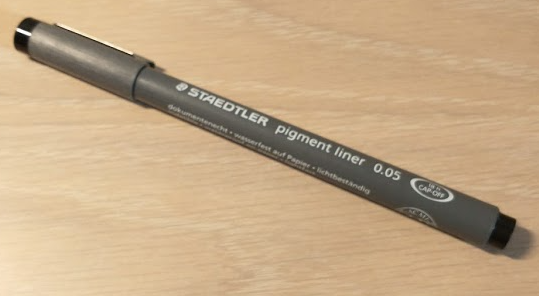

15. 90-45-45 set square

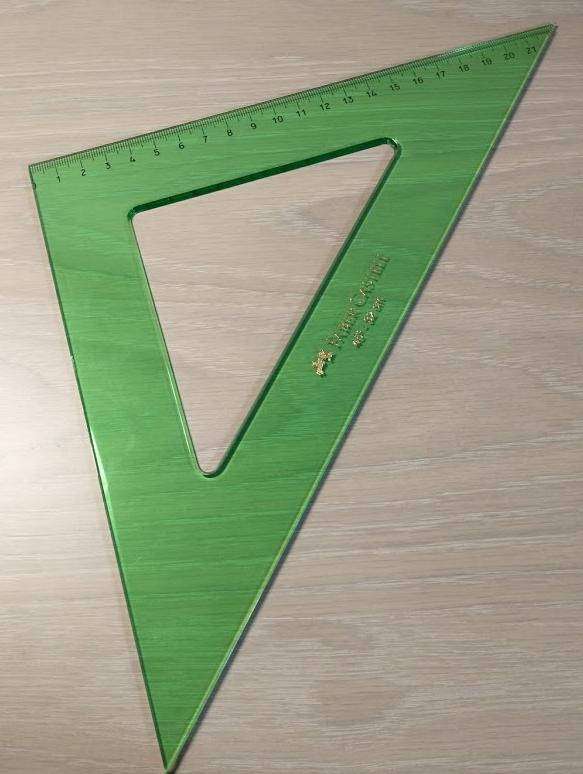

16. Seeds

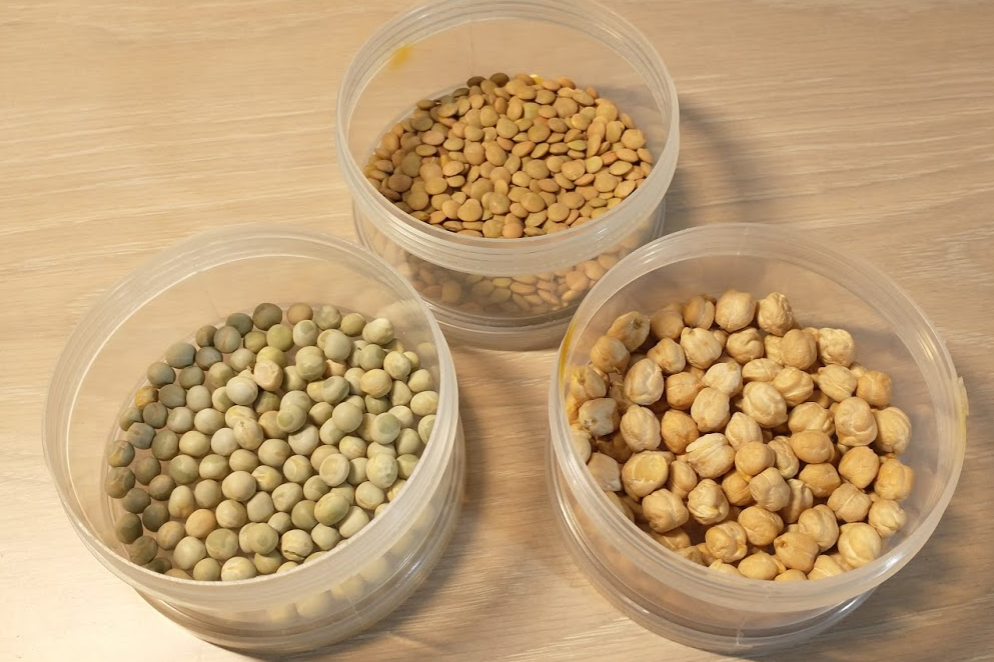

Instructions:

Step 1: Attach the Velcro to the cardboard to close one side of the box

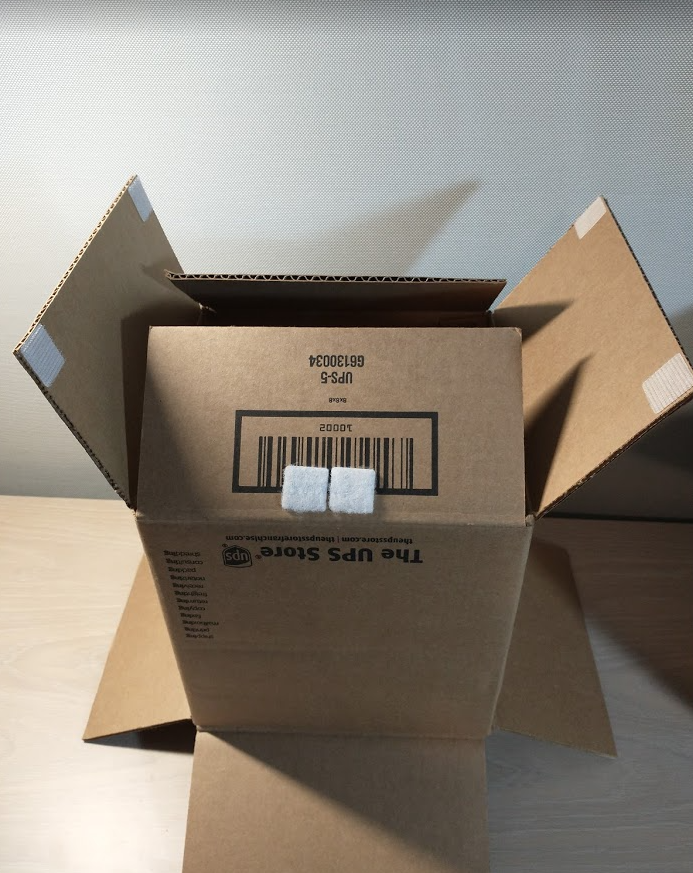

Step 2: Cut a piece of the cardboard to the size of the lid and paste it with glue

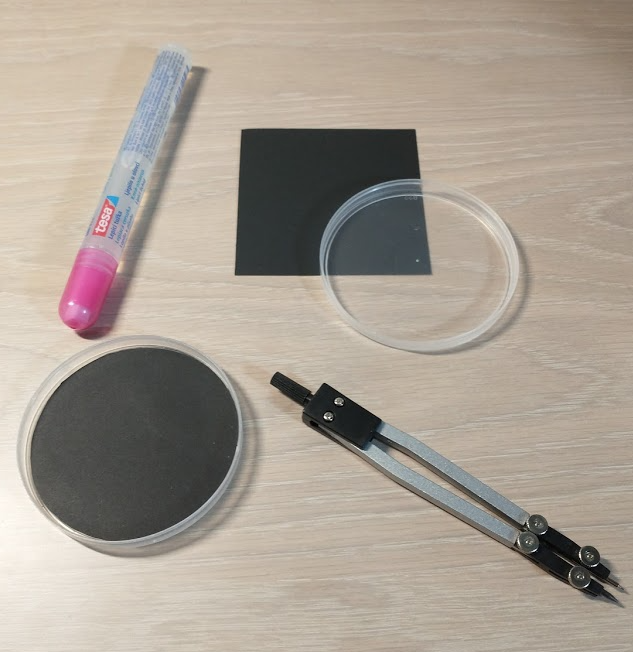

Step 3: Cut 3 pieces of aluminum foil to the size of the box and attach it to 3 sides of the box with glue

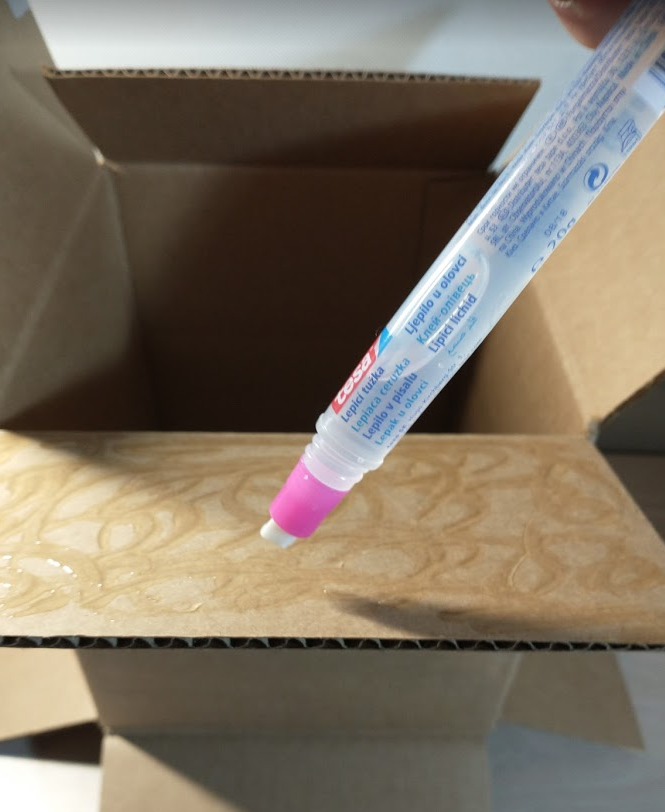

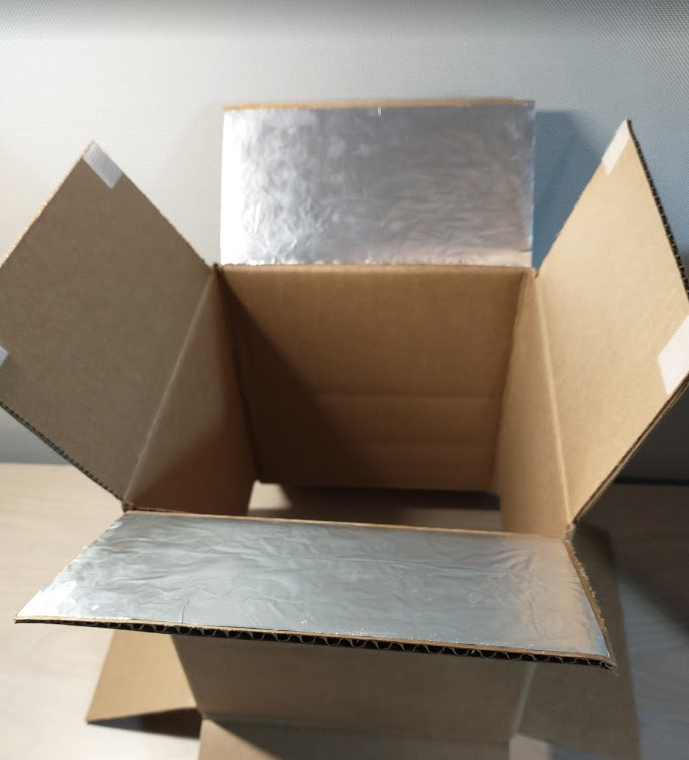

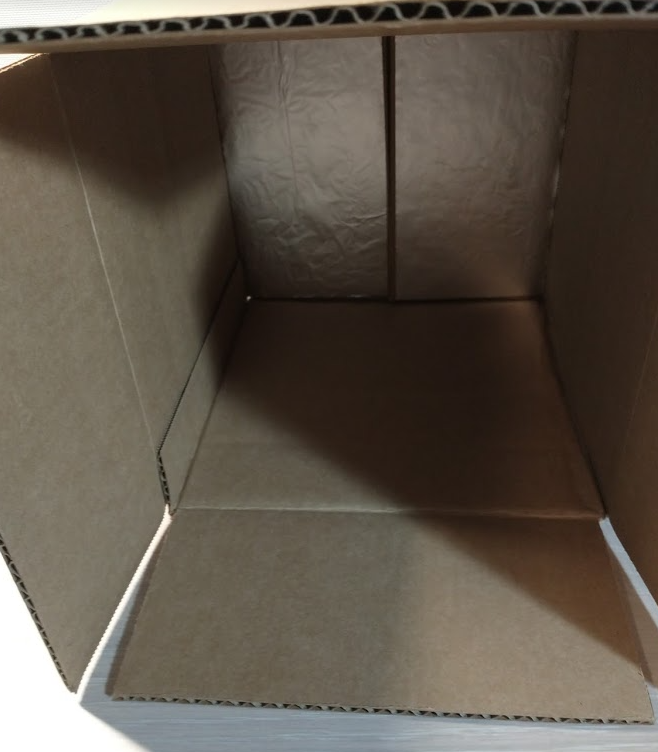

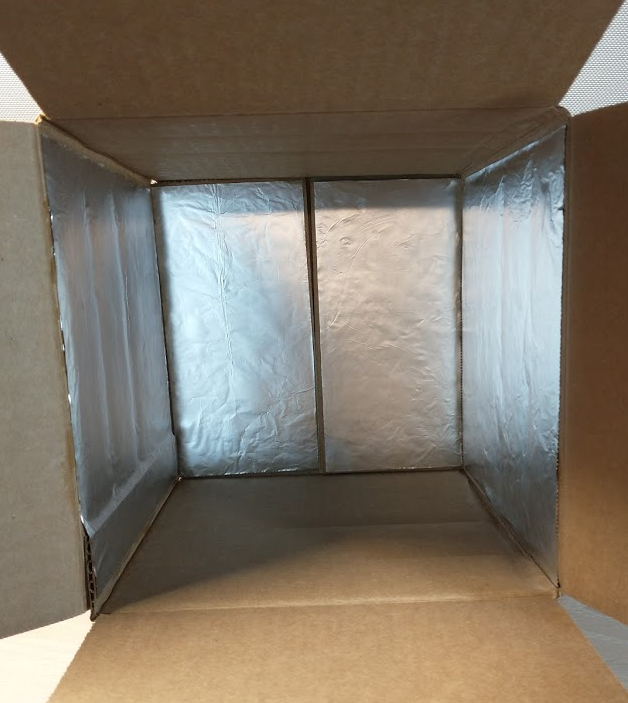

Step 4: Cut a piece of cardboard to match the size of the box. Then, draw two crosses: one measuring 8 cm x 8 cm and another that's half that size. There must be a distance of 9cm between the center of the larger cross and the right horizontal edge of the small cross.Also, draw a line parallel to the left horizontal edge of the larger cross 9 cm apart.

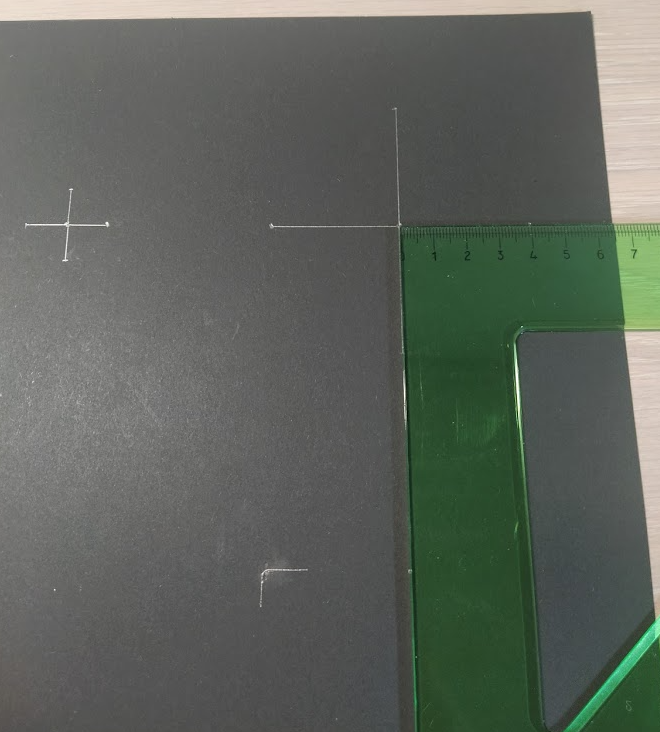

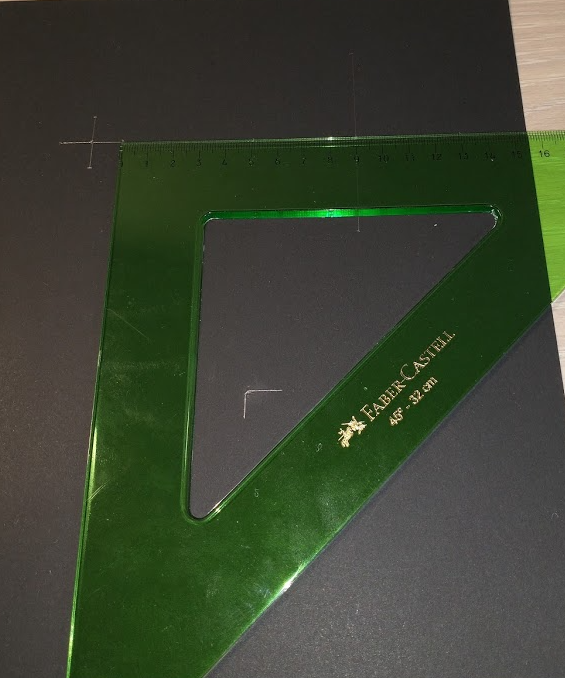

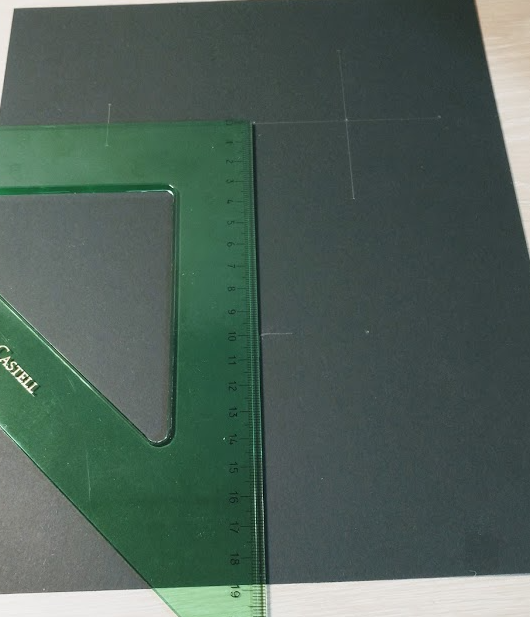

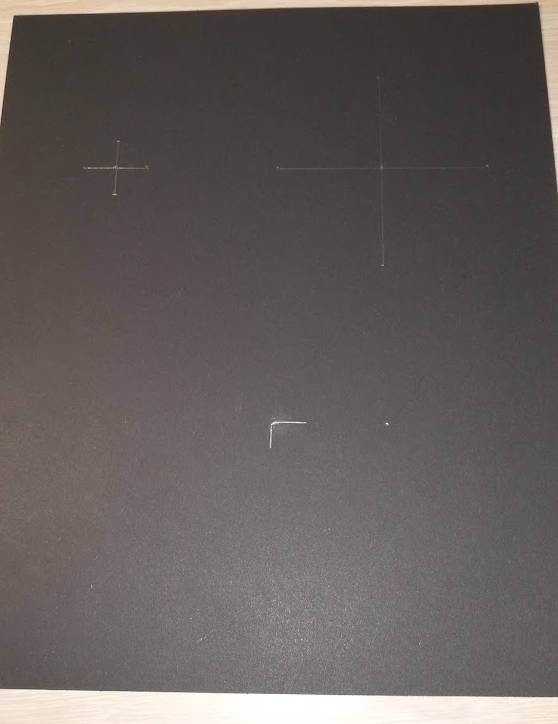

Step 5: Attach the cardboard to 1 sides of the box with glue

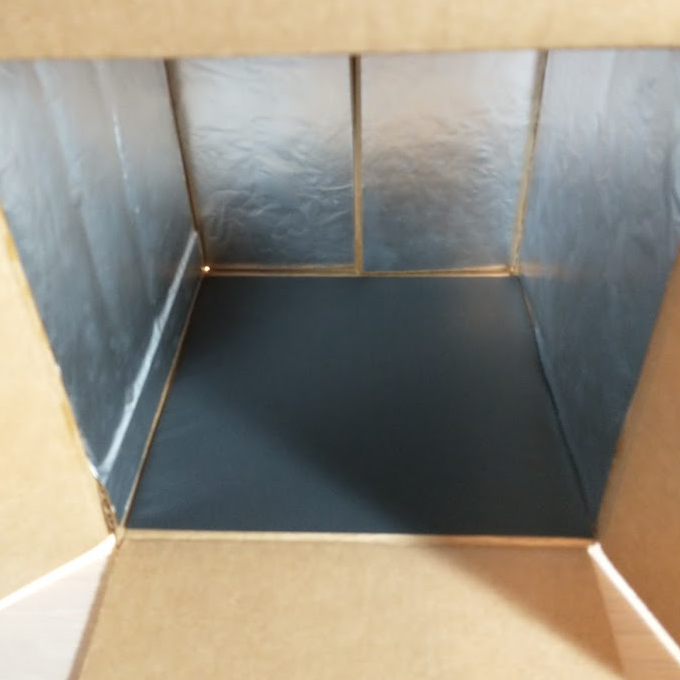

Step 6: Draw a rectangular hole in the middle of the box. Make sure that the hole is 6 cm and 7 cm away from the box's edges

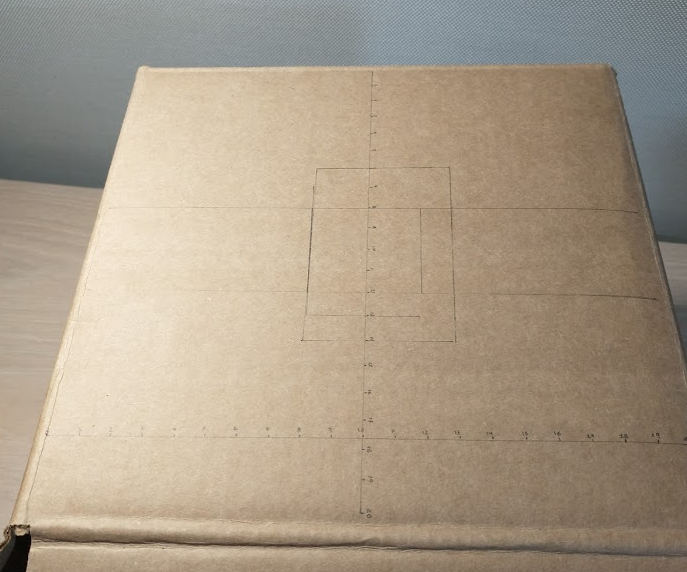

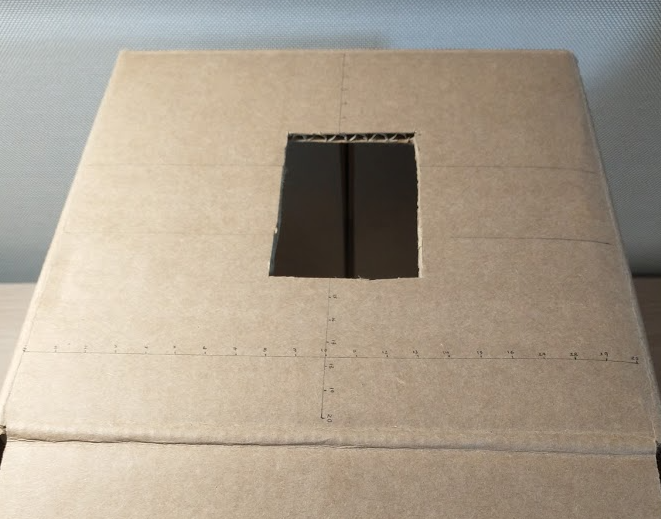

Step 6: Attach the LED light inside the box, adjacent to the hole

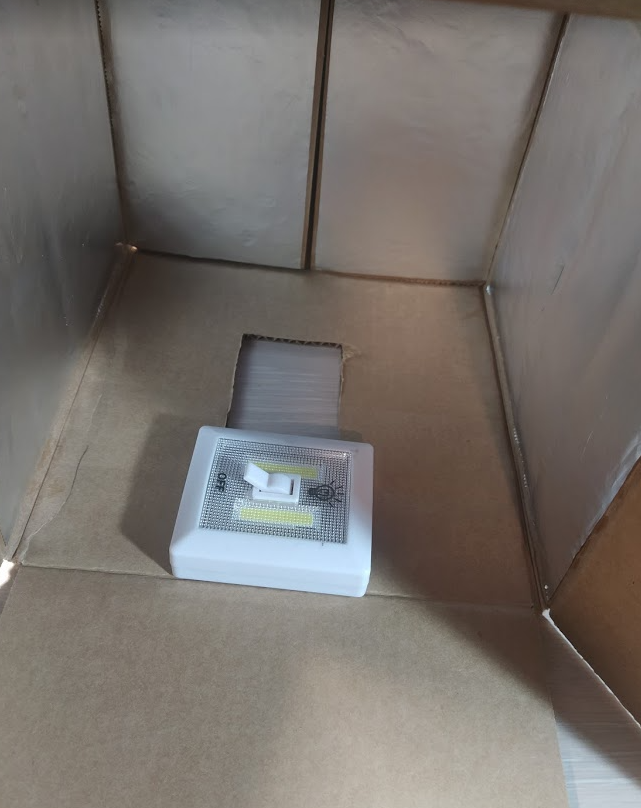

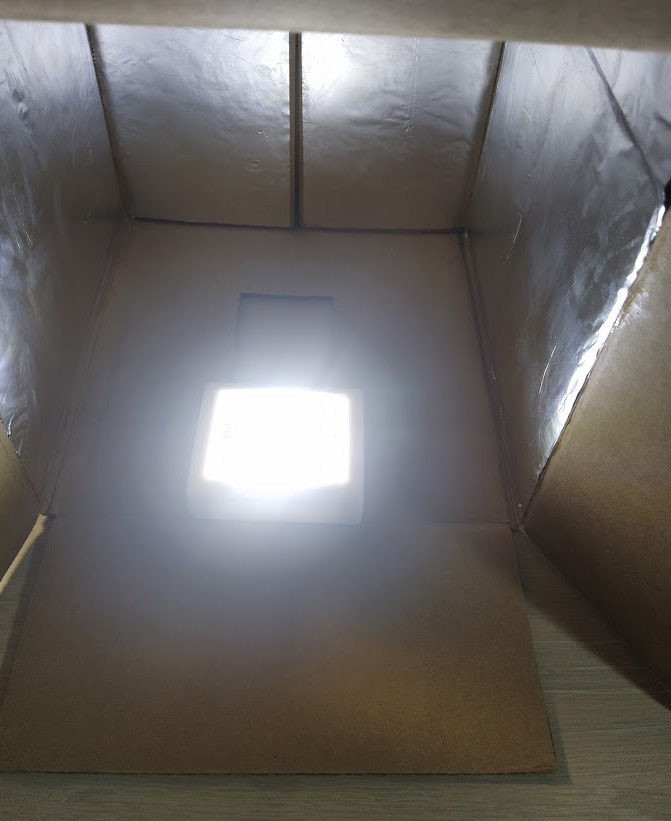

Step 7: Place the cellphone on top of the box

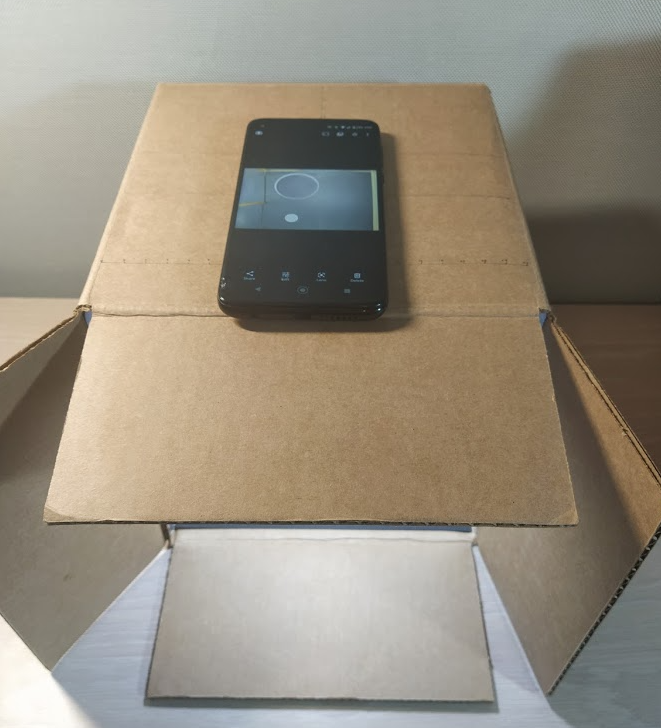

Step 7: Place the seeds, coin and envelope inside the box

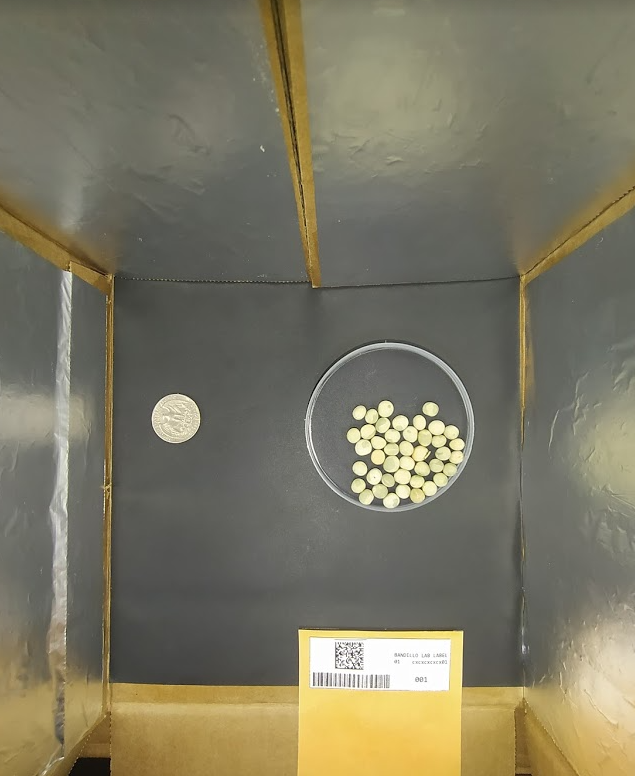

**Appendix 2**

Ground Truth Verification Seed Size

A linear trend model is computed for Seed Size Rep 1 (mm) given the Average of Major Axis Length (Shape) (mm). The model may be significant at p <= 0.05.

Model formula:	( Average of Major Axis Length (Shape) (mm) + intercept )

Number of modeled observations:	61

Number of filtered observations:	0

Model degrees of freedom:	2

Residual degrees of freedom (DF):	59

SSE (sum squared error):	9.0803

MSE (mean squared error):	0.153903

R-Squared:	**0.872852**

Standard error:	0.392305

p-value (significance):	< 0.0001

A linear trend model is computed for Seed Size Rep 2 (mm) given the Average of Major Axis Length (Shape) (mm). The model may be significant at p <= 0.05.

Model formula:	( Average of Major Axis Length (Shape) (mm) + intercept )

Number of modeled observations:	61

Number of filtered observations:	0

Model degrees of freedom:	2

Residual degrees of freedom (DF):	59

SSE (sum squared error):	11.6753

MSE (mean squared error):	0.197886

R-Squared:	**0.848112**

Standard error:	0.444843

p-value (significance):	< 0.0001

A linear trend model is computed for Seed Size Rep 1 (mm) given the Average of Minor Axis Length (Shape) (mm). The model may be significant at p <= 0.05.

Model formula:	( Average of Minor Axis Length (Shape) (mm) + intercept )

Number of modeled observations:	61

Number of filtered observations:	0

Model degrees of freedom:	2

Residual degrees of freedom (DF):	59

SSE (sum squared error):	9.84116

MSE (mean squared error):	0.166799

R-Squared:	**0.862198**

Standard error:	0.408411

p-value (significance):	< 0.0001

A linear trend model is computed for Seed Size Rep 2 (mm) given the Average of Minor Axis Length (Shape) (mm). The model may be significant at p <= 0.05.

Model formula:	( Average of Minor Axis Length (Shape) (mm) + intercept )

Number of modeled observations:	61

Number of filtered observations:	0

Model degrees of freedom:	2

Residual degrees of freedom (DF):	59

SSE (sum squared error):	11.5097

MSE (mean squared error):	0.19508

R-Squared:	**0.850265**

Standard error:	0.441679

p-value (significance):	< 0.0001

Individual Regression lines:

Panes	Line	Coefficients

Row	Column	p-value	DF	Term	Value	StdErr	t-value	p-value

Seed Size Rep 1 (mm)	Average of Major Axis Length (Shape) (mm)	< 0.0001	59	Average of Major Axis Length (Shape) (mm)	**0.841977**	0.0418369	20.1253	< 0.0001

intercept	0.456423	0.322714	1.41433	0.162522

Seed Size Rep 1 (mm)	Average of Minor Axis Length (Shape) (mm)	< 0.0001	59	Average of Minor Axis Length (Shape) (mm)	**0.829191**	0.0431572	19.2133	< 0.0001

intercept	0.563024	0.332501	1.6933	0.0956728

Seed Size Rep 2 (mm)	Average of Major Axis Length (Shape) (mm)	< 0.0001	59	Average of Major Axis Length (Shape) (mm)	**0.861058**	0.0474397	18.1506	< 0.0001

intercept	0.325294	0.365932	0.888947	0.377642

Seed Size Rep 2 (mm)	Average of Minor Axis Length (Shape) (mm)	< 0.0001	59	Average of Minor Axis Length (Shape) (mm)	**0.854288**	0.0466726	18.3038	< 0.0001

intercept	0.386334	0.359586	1.07439	0.287024

**Appendix 3**

Correlation Values

Seed 1 and Seed 2	**0.896596**

Avg (Seed 1,Seed 2) Vs Minor Axis Length **0.950142**

Avg (Seed 1,Seed 2) Vs Major Axis Length **0.952414**
